# The cytosolic PRMT5-CDK4 complex impairs cell cycle kinase signaling

**DOI:** 10.64898/2025.12.11.693714

**Authors:** Sarah Masser, Tanja Lotteritsch, Anton Emil Ludwig, Bettina Halwachs, Elisabeth Annerer, Marion Mussbacher, Ulrich Stelzl

## Abstract

Both, protein arginine methyltransferases (PRMTs) and protein kinases are critical regulators of cellular processes and frequently dysregulated in malignancy. To systematically study the crosstalk of the two regulatory enzyme classes, we used parallel yeast two-hybrid matrix screening and defined 45 interactions connecting 4 PRMTs and 20 human kinases. The PRMT–kinase network revealed a strong association between PRMTs and cell cycle/mitogen-activated kinases. Notably, the PRMT5–CDK4 emerged as the most prominent functional cell cycle link through integrative data analyses of different high-throughput datasets. Mechanistic cell biological studies confirmed that the PRMT5–CDK4 complex localizes exclusively in the cytosol, where its formation was enhanced during a G1/S cell cycle block. This PRMT5 interaction modulated the CDK4–CCND3 and CDK4–CDKN2A interaction dynamics, irrespective of PRMT5’s methyltransferase activity. Phospho-proteomic profiling demonstrated that PRMT5 overexpression functionally phenocopies the signaling effects of pharmacological CDK4 inhibition with Palbociclib. The findings establish a non-enzymatic regulatory role for PRMT5, proposing it functions as an inhibitory modulator of CDK4-driven oncogenic signaling.

## INTRODUCTION

Protein-protein interactions (PPI) and post-translational modifications (PTM) - such as phosphorylation and arginine methylation (RMe) - are fundamental molecular factors driving cancer progression due to their broad impact on signaling networks and cellular homeostasis^1–4^. Dysregulation of signaling cascades, particularly within the core cell cycle machinery, contributes to aberrant cell division, a hallmark of cancer and a central force in tumorigenesis. Thus, several anti-cancer strategies have focused on halting tumor progression by targeting critical regulators of the cell cycle. These include clinically advanced kinase inhibitors targeting AURKA (Alisertib), CDK4/6 (Palbociclib, Abemaciclib), CHK1 (Prexasertib), and PLK1 (Volasertib) (*reviewed in^5–7^)*. Mammalian protein arginine methyltransferases (PRMTs), which are ubiquitously expressed^8, 9^, transfer methylation marks on target proteins. PRMTs 1, 4, 6, and 8 catalyze asymmetric R–dimethylation (type I, ADMA) while PRMT5 - recognized as the major type II PRMT enzyme - either monomethylates or symmetrically dimethylates arginine side chains (SDMA) of histone and non-histone proteins^9^. Thereby, PRMTs regulate influential cellular processes such as chromatin remodeling, gene transcription, and cellular differentiation. PRMT5 in particular also plays a fundamental role in pre-mRNA splicing and RNA processing^10^. PRMT5 functions in the methylosome, a hetero-octameric complex including the essential PRMT5 coactivator WDR77 ^11, 12^ and other flexibly tethered substrate adaptors^13–15^, and has emerged as a pivotal target in cancer research.

The combined loss of *MTAP* with the tumor suppressor *CDKN2A* through co-deletion (on chr9:21, GRCh38 ∼100kb apart) represents the most prevalent chromosomal deletion events across cancers^16^. *CDKN2A* encodes for p16/INK4a protein (here CDKN2A), the most important inhibitor of CDK4/6 in the cell cycle^17^. MTAP protein is a methylthioadenosine phosphorylase responsible for the first step in the methionine salvage pathway acting after MTA (5’-Methylthioadenosin) has been generated from S-adenosylmethionine. Mechanistically, *MTAP*^null^ tumors, accumulate the MTA metabolite which selectively impairs PRMT5 activity, rendering these cancers more susceptible to PRMT5 inhibition (PRMT5i), creating synthetic lethal vulnerability^16, 18^, which is leveraged in cancer therapy. Therefore, a new class of MTA-cooperative PRMT5i (MTAC-PRMT5i including e.g. MRTX1719) has emerged^19–21^. These inhibitors are demonstrating high selectivity and efficacy in the clinic, mitigating non-selective toxic effects observed in non-MTAC-PRMT5i^22, 23^. Furthermore, combining CDK4/6i and PRMT5i is a promising strategy for treating cancers, especially those with *CDKN2A* loss and high CDK4/6 activity, due to shared roles in governing the G1 checkpoint and downstream effects via the PRMT5–MDM4–p53 axis^24, 25^. Accordingly, MTAC-PRMT5i are currently undergoing clinical evaluation in *MTAP*^null^ cancer as combination therapy with CDK4/6i and other cell cycle–targeting agents (NCT06810544, NCT06333951, NCT05094336, NCT04794699).

A number of additional reports suggest that PRMT–kinase interplay strongly impacts cellular signaling^26–30^, while on the other hand no protein methyltransferase-kinase interactions were detected in large reference AP-MS or CF-MS studies^3^. This led us to systematically investigate the interaction space between five human PRMTs and a large panel of 180 human protein kinases using a matrix yeast-two-hybrid (Y2H) approach. We established a PRMT–kinase interaction network with 45 interactions involving 20 kinases. A highly significant interactome overlap of PRMT5 and CDK4 supported by integrative data analysis, including the use of independent data from native organelle IPs^31^, highlights PRMT5 as regulator of the cell cycle. High scoring AlphaFold multimer predictions^32^ of a hetero-multi(do-/deca)meric CDK4–PRMT5 complex shows the potential of PRMT5 to directly regulate CDK4 activity. Our mechanistic cell biological analyses focused on the interaction between PRMT5 and CDK4. In live-cell microscopy imaging, the PRMT5–CDK4 complexes were exclusively cytoplasmic and this interaction increased during G1/S cell cycle arrest. Quantitative co-IP PPI assays^33, 34^ demonstrated that, independent of its methyltransferase activity, PRMT5 binding modulates CDK4 activity through enhancing complex formation of CDK4 with its key regulators, CDKN2A and CCND3. This suggested a regulatory role tuning subcellular dynamics and composition of CDK4 complexes. Mass spectrometry (MS)-based phospho-proteomic analysis demonstrated that PRMT5 overexpression phenocopies the signaling effects of pharmacological CDK4/6i. Our study suggests that enhanced, cytosolic PRMT5–CDK4 interaction has an inhibitory effect on cell cycle progression.

## RESULTS

### A PRMT–kinase interaction network links PRMTs to cell cycle regulation

We utilized a Y2H matrix approach to systematically identify binary interactions between five human PRMTs and 180 human kinases. The Y2H approach offers a major methodological advantage in detecting transient and less stable interactions, particularly relevant for kinases. Genetic reporter gene activation amplifies interaction signals in a binding equilibrium also at very low protein expression levels^35–38^. 170 active protein kinases and 10 non-catalytically kinase subunits such as cyclins (CCNs) or PRKRA were prioritized based on activity measurements in yeast^39, 40^. Covered by 200 individual ORFs, this human protein kinase set is a representative subset of the human kinome including members of all major protein kinase families^41^. As potential interaction partners, we selected 5 PRMTs covered by 9 individual ORFs, including type I PRMTs (PRMT1, 6, 8, and CARM1/PRMT4) and the key type II PRMT5. To enhance sensitivity of the Y2H screen while maintaining specificity^42–44^, we tested each protein pair in eight configurations (C- and N-terminal vector fusions and bidirectional orientations) in a pairwise all-against-all Y2H matrix approach. We tested close to 17,000 PPIs by pairing 368 prey/377 bait kinase constructs with seven bait/eight prey PRMT constructs in duplicate in a 384-format matrix screen (prey-2-bait / bait-2-prey) setup^45^ (**Figure 1A**). We identified 870 raw hits by colony growth, from which we excluded 16 potentially auto-active kinase constructs and weak interactions which appeared only once. As a result, 104 bait-prey pairs, identified in duplicates, were considered as high-confident interactions and included for further evaluation (**Figure 1B, Suppl. Table1**, representative agar shown in **Figure S1A**). The resulting high-confidence PRMT–kinase network was visualized with growth counts and vector configuration of the bidirectional multi-tagging approach (**Figure 1C**). Interaction frequency and strength varied by PRMT–kinase pair and vector setup, producing a spectrum of Y2H counts for interacting protein pairs (**Figure 1B-C**). Due to relatively low Y2H counts and the stringent cut-off, we formally excluded all PRMT1 pairs. However, given the ∼80-85% sequence identity between PRMT1 and PRMT8^46^, the interactions likely reflect overlapping partners. Collapsing of the 104 high-confidence PRMT-kinase Y2H interactions finally resulted in 45 unique PRMT–kinase interaction pairs linking 4 PRMTs with 20 human protein kinases (**Figure 1C, right, Suppl. Table2**). Notably, PRMT interactions showed a preference for binding of CMGC kinases (including MAPKs, CDKs, and CDKLs), which comprised approximately 30% of the identified PPIs, followed by CAMK family members at around 25% (**Figure 1D**). PRMT5 and PRMT6 exhibited the highest number of interactions with cell cycle related kinases (CDKs, AURKA/B, and PLK1), highlighting directly links of these PRMTs to cell cycle regulatory kinases. To explore the kinase-related biological roles in more detail, high-confidence interactions of individual PRMTs were analyzed for kinase-driven reactome pathway enrichment. Although some of the enriched pathway terms were rather generic for kinases and shared between the methyltransferases, e.g., nuclear events (kinase and TF-activation), selectively enriched pathways were evident. While the kinases interacting with CARM1, which were primarily associated with a MAPK-ERK signature (**Figure S1B**), PRMT5 (**Figure 1E**) and PRMT6 (**Figure S1C**) showed significant enrichment in cell cycle control, transcription regulation, and stress/damage-induced cellular responses. In line with reported evidence^47, 48^, PRMT6 kinase partners were involved in senescence-related processes, whereas PRMT5 was more strongly linked to cell cycle-associated kinases in the enrichment analysis. In summary, we report 45 binary protein interactions (from 104 high-confidence Y2H pairs) between four PRMTs and 20 human kinases, preferentially from the CDK and MAPK kinase families.

**Figure 1:**
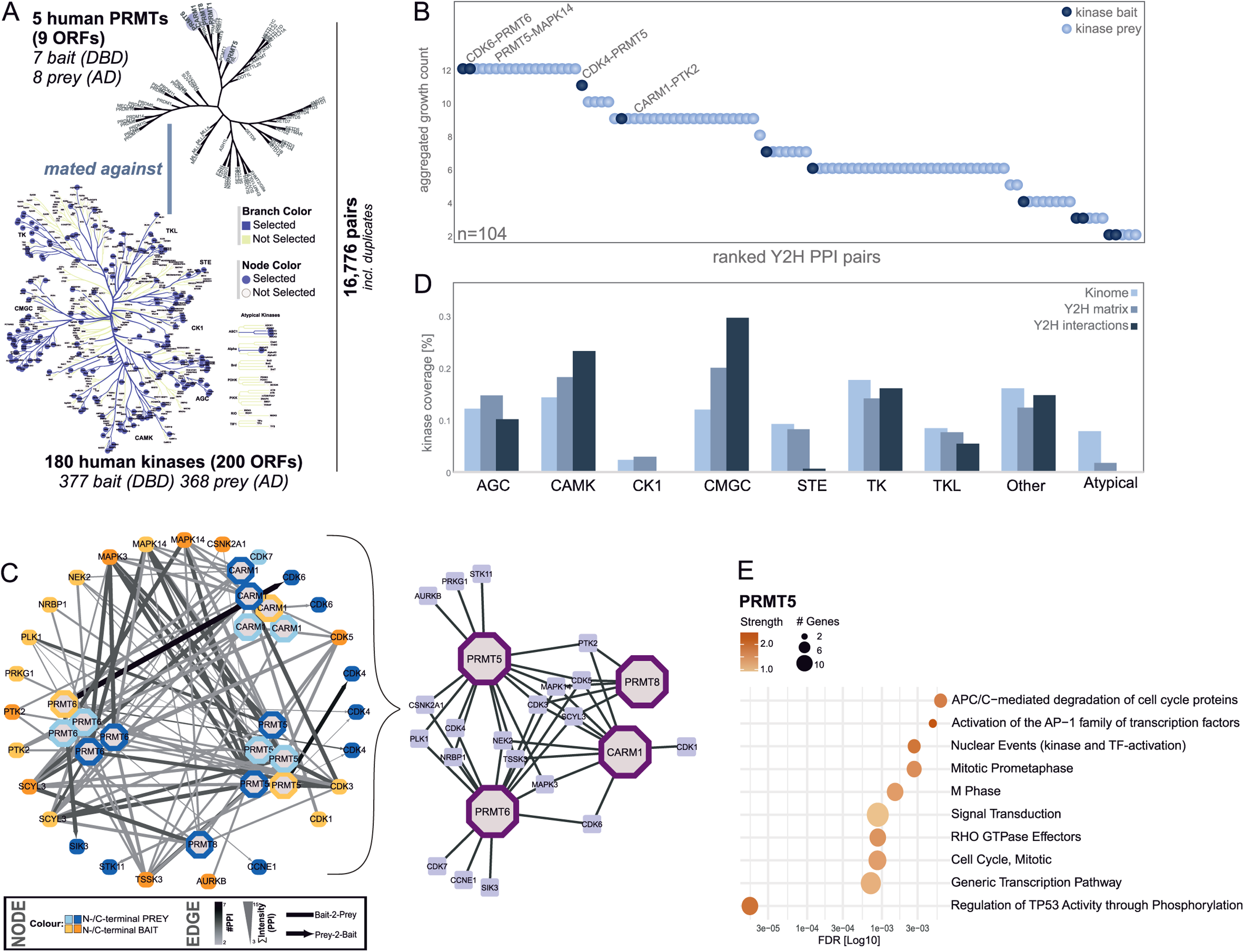
PRMT–Kinase protein interaction network. **(A)** Schematic overview of active human kinases (*sourced from KinHub*) and PRMTs (*sourced from ChromoHub*) used in a yeast two-hybrid (Y2H) interaction screen. Kinases and PRMTs are shown on their phylogenetic trees to illustrate sequence relationships. 180 human kinases (200 ORFs) are represented by 377 bait (DBD) and 368 prey (AD) constructs. 5 human PRMTs (9 ORFs) are represented by 7 bait (DBD) 8 prey (AD) for pairwise testing interactions in a pairwise all-against-all Y2H matrix approach. **(B)** Ranked dotplot showing 104 prey–bait pairs identified by Y2H screening that appeared in duplicate with the total colony growth count for each kinase-PRMT construct pair (prey kinase constructs dark blue, bait kinase constructs light blue). **(C)** PRMT-kinase interaction network. left: Network representation of 104 PRMT–kinase pairs, with edges indicating interaction frequency (color) and ∑growth counts (PPI strength; width). Node colors denote the N- or C-terminal fusion proteins. Right: final, collapsed PRMT–Kinase interaction network with 45 interactions between 4 PRMTs and 20 Kinases. **(D)** Bar plot highlighting the fraction of interacting kinases relative to the input Y2H matrix and to the overall kinome, shown per kinase family. (**E**) Functional analysis of uniquely identified PPIs across all vector configurations for PRMT5, highlighting the top 10 most enriched kinase-driven Reactome pathways, ranked by FDR scores obtained from the String-DB analysis.

### Functional and structural evidence for a multimeric PRMT5-CDK4 complex

Both PRMTs and kinases broadly regulate key cellular processes within complex networks. Therefore, we hypothesized that analyzing their interactome could reveal functional connections through their interaction partners. We leveraged the person correlation-based RNA-seq gene-gene co-expression matrix from Enrichr^49^ to assess the top 100 genes most strongly co-expressed with PRMT5 (**Suppl. Table3**). The PRMT5 gene set was then compared to the top 300 genes linked to each kinase in the ARCHS4 dataset to evaluate their similarity. This co-expression gene analysis revealed that PRMT5 and CDK4 exhibited the most significant similarity (odds ratio >160, p-value = 9.64e^-96^) among all kinase sets (**Figure 2A, S2A**). Other kinases, which we also identified as PRMT5 interactors in the Y2H screen including PLK1 and AURKB, were within the top 10% of ranked kinases with PRMT5 (odds ratio: >10 - <120) (**Figure 2B**). In agreement with the known PRMT5 biology, RIOK1, a well-established co-complex member of the methylosome^50^, was also identified among the most significantly overlapping correlation profiles in this analysis. The convergence of kinases identified through gene set co-expression analysis and Y2H interaction screening highlights their involvement in cell cycle–related processes. This can be illustrated by a word cloud generated from String-DB^51^ enrichment analysis of overlapping interaction partners from the PRMT5 interactome and PRMT5-Y2H kinase screen, showcasing the most significantly enriched functional terms (Reactome, Keywords, GO color-coded by the FDR values; **Figure 2C**). It is further supported by the Reactome-based pathway enrichment analysis of the top 100 co-expressed genes for PRMT5 and CDK4, respectively, which in both cases showed predominant enrichment in cell cycle–associated pathways (**Figure 2D**). As CDK4 is one of the top interactions in the Y2H analysis and emerged as the functionally most similar PRMT5 partner in the annotation analysis, the PRMT5–CDK4 interaction was prioritized for further analyses.

**Figure 2:**
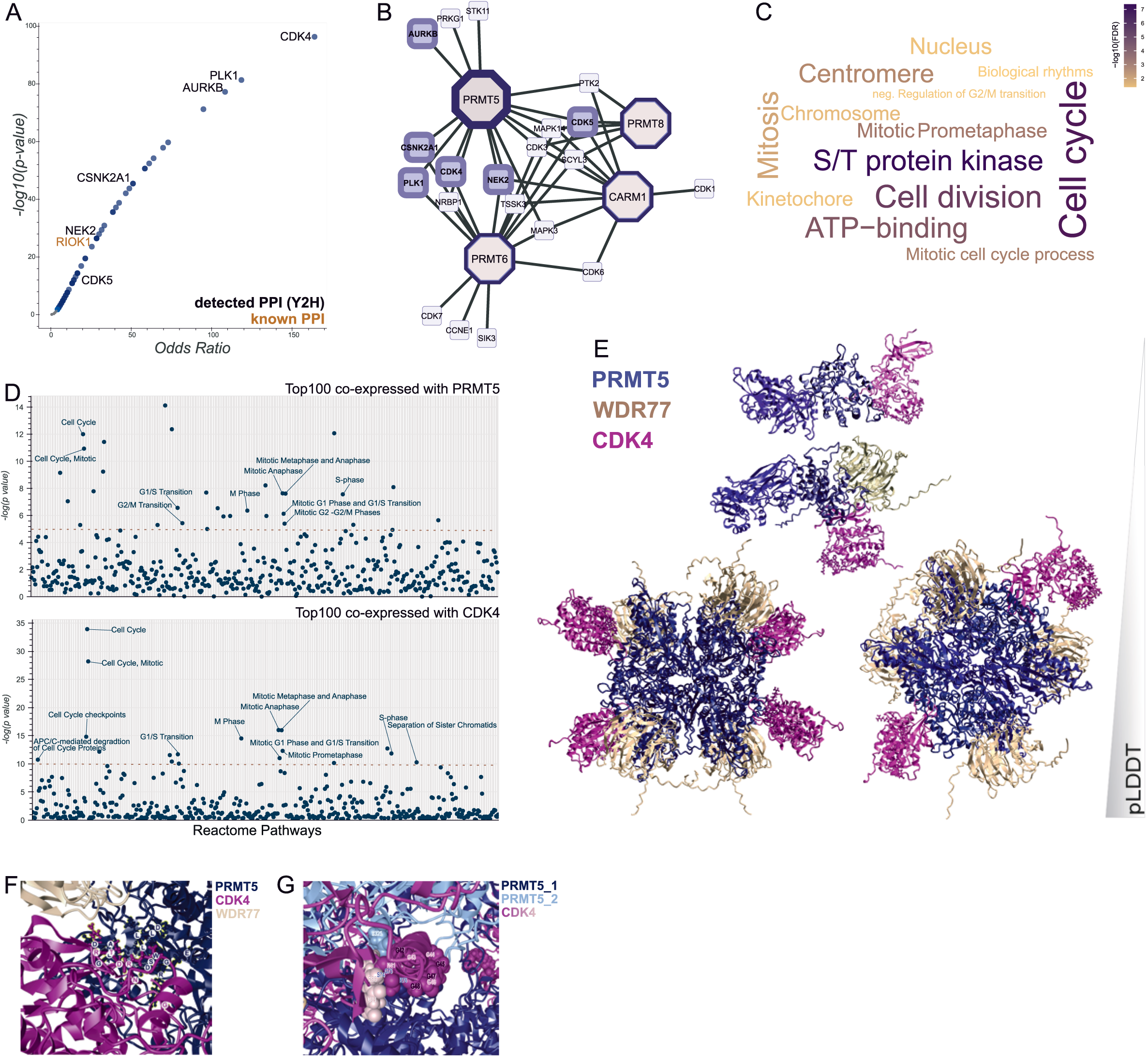
Data integration reveals the importance of a hetero-multimeric PRMT5-CDK4 complex in cell cycle regulation. (**A**) Scatterplot displaying all kinases with significant co-expression patterns with PRMT5 (top 100 genes). Gene–gene correlation analysis using ARCHS4 RNA-seq data was performed with Enrichr. The x-axis shows the odds ratio, and the y-axis the significance as -log₁₀(p-value). Kinases identified in the Y2H screen are labeled in black, RIOK1, a known PRMT5 kinase interactor is highlighted in brown. (**B**) Network representation of PRMT5 and its top co-expressed kinases, illustrating the overlap between the co-expression analysis data and the established binary interaction data (c.f. Fig. 1C). (**C**) String-DB enrichment analysis of overlapping kinases from the PRMT5 interactome (A) and PRMT5-Y2H kinase screen (Figure 1) The top enriched functional terms (Reactome, Keywords, GO) from the intersection of co-expressed and interacting kinases were visualized as a word cloud with significance values color coded. (**D**) Manhattan plots showing Reactome pathway enrichment of the top 100 co-expressed genes with PRMT5 (top) and CDK4 (bottom). Notably, both profiles highlight significant enrichment of cell cycle–related pathways. (**E**) Structural prediction of the PRMT5–CDK4 interaction, modeled with the coactivator WDR77 using AlphaFold-Multimer. Predictions were generated for increasing levels of structural complexity: starting from the binary PRMT5 (blue)–CDK4 (purple) complex (pLDDT = 0.48), progressing to the symmetric hetero-dodecameric PRMT5–WDR77 (gold)–CDK4 complex (pLDDT = 0.84–0.87). (**F-G**) Zoom in to the predicted interaction interface between PRMT5 and CDK4 in the AlphaFold-Multimer hetero-dodecameric model (**F**), highlighting interacting residues in a glycine-rich motif N_41_GGGGGG_48_(**G**).

Next, we applied AlphaFold Multimer^32^ to predict the PRMT5–CDK4 complex structure. AlphaFold Multimer has been benchmarked by others as highly predictive of interaction interfaces^52^. We first modeled the monomeric PRMT5–CDK4 complex (pLDDT = 0.48), and improved prediction accuracy by including the most important PRMT5 cofactor, WDR77, in a PRMT5–WDR77–CDK4 complex (pLDDT = 0.58; **Figure 2E, top**). To mimic the native assembly, we next modeled symmetric hetero-octamers of PRMT5–WDR77 with two or four CDK4 units, yielding high-confidence structures with bound CDK4 with pLDDT scores of 0.87 for the decamer and 0.84 for the dodecamer (**Figure 2E, bottom**). While the very high pLDDT score may partly stem from the publicly available methylosome structure (4GQB^12^) the models support a PRMT5–CDK4 interaction within the a hetero-do-/decameric complex.

To interpret our results in more detail, we superimposed two predicted PRMT5–WDR77–CDK4 monomers onto the original cryo-EM structure of the core methylosome at an RMSD of 0.7567Å (**Figure S2B**). This allowed us to analyze the interface contacts within the predicted dodecamer comparing both superimposed (**Figure S2B,C**) and predicted dodecameric structures (**Figure 2E-F, S2D**). PRMT5 engages CDK4 via its N-terminal TIM barrel domain (AA 13–292), specifically through A27–Q36, and in the predicted complex via an interface spanning D70–D77. This second interaction site involves CDK4 residues D84 and R85, the latter forming a strong ionic interaction with E325 of the adjacent PRMT5 Rossmann fold (**Figure 2E, S2D**). From the CDK4 perspective, key residues involved in the PRMT5 interaction define a distinct interface at the N-lobe of the kinase fold (**Figure S2C-D**), including the amino acids E11, G13, V14, and Y17 (**Figure S2C**) within and around the conserved glycine-rich loop (G-X-G-X-X-G), which is critical for ATP binding, as well as CDC37, HSP90, and CCND binding^53^. Furthermore, another glycine-rich motif in the N-lobe comprising the residues N41, G43, G44, and G46 (**Figure 2G**) formed contacts with PRMT5 in the multimeric complex. Finally, contacts were also seen at the N-lobe–hinge junction with R82, T83, D84, R85, and K88 (**Figure S2C-D**). The CDK4 N-lobe region dictates complex stability, kinase conformation and binding specificity, especially CCND1/3 and CDKN2A binding that regulates CDK4 activity^17, 54^. The ensemble of predicted PRMT5–CDK4 interaction sites suggests that PRMT5 binding may modulate the binding of activators or inhibitors, as the involved residues are located in a bridge between the canonical effector binding interfaces (see below). Collectively, we modeled a high-scoring hetero-multimeric PRMT5–WDR77–CDK4 complex, revealing a large interface between PRMT5 and CDK4, which possibly enables allosteric modulation of CDK4 function through its key regulators CCND1/3 and CDKN2A.

### PRMT5 interacts with CDK4 in the cytosol in a cell cycle-dependent manner

To monitor the PRMT5–CDK4 interaction in living cells and capture potential subcellular dynamics, we traced the PPI through live cell confocal fluorescence microscopy and cell counting, employing a Venus-EYFP split complementation system^55–57^ (**Figure 3A**). The small(F1) and large(F2) Venus-fusion tags were systematically appended to the C-termini of PRMT5 and CDK4 in both configurations, PRMT5_F1_–CDK4_F2_ and CDK4_F1_–PRMT5_F2_ minimizing potential tagging bias. We transiently co-transfected complementation construct pairs into HEK293T cells, where EYFP fluorescence visualizes the direct PPI and their specific subcellular localization, independent of other cellular regions where either protein may localize individually. While specificity tests pairing our PRMT5 or CDK4 constructs with other complementation constructs did not support the folding of EYFP (e.g. WDR77–CDK4, **Figure S3A**), the interaction between PRMT5 and CDK4 exhibited a strong complementation signal in both configurations. The cells were subjected to cell cycle synchronization treatments, followed by qualitative and quantitative fluorescence image analysis. Interestingly, while the EYFP fluorescence signal, indicative of the PRMT5–CDK4 interaction, was observed in unsynchronized, starved (G0) and double thymidine blocked cells (G1/S), comparison across conditions showed a distinct, evenly cytosolic localization of the complex with enhanced signal intensity in the cells arrested in G1/S phase (**Figure 3B**). To quantify overall PPI levels in cells arrested in the different cell cycle phases, we measured EYFP complementation signal intensity using flow cytometry (**Figure S3A**) and quantitatively assessed our confocal images (**Figure S3B**). Across three independent experiments, each evaluating at least 600 cells per condition, we observed a marked increase of the PRMT5–CDK4 interaction during the G1/S phase compared to unperturbed cells (**Figure 3C, S3C**). In contrast, the EYFP complementation signal of the PRMT5–PRMT5 homomeric interaction did not show qualitative (**Figure 3B**) nor quantitative (**Figure 3C, S3C**) cell cycle-dependent differences. We performed the same set of experiments with an enzymatically inactive PRMT5^null^ (G367A/R368A)^58^ mutant version with very similar outcome (**Figure 3D**). We observed the same increase in cytosolic localization in the G1/S treated cells of the PRMT5^null^–CDK4 interaction and therefore concluded that the interaction is independent of the methyltransferase activity of PRMT5. Although both proteins are expressed in the cytosol and nucleus (PRMT5 in cytosol and nucleoplasm; CDK4 in nucleoplasm, cytosol, nuclear membrane, and nucleoli^59^). Using EYFP complementation as readout, the interaction was observed as uniformly distributed signal in the cytosol, and we did not detect any interaction with nuclear localization. This suggests that under the tested conditions CDK4–PRMT5 complexes do not shuttle from the cytosol to the nucleus.

**Figure 3:**
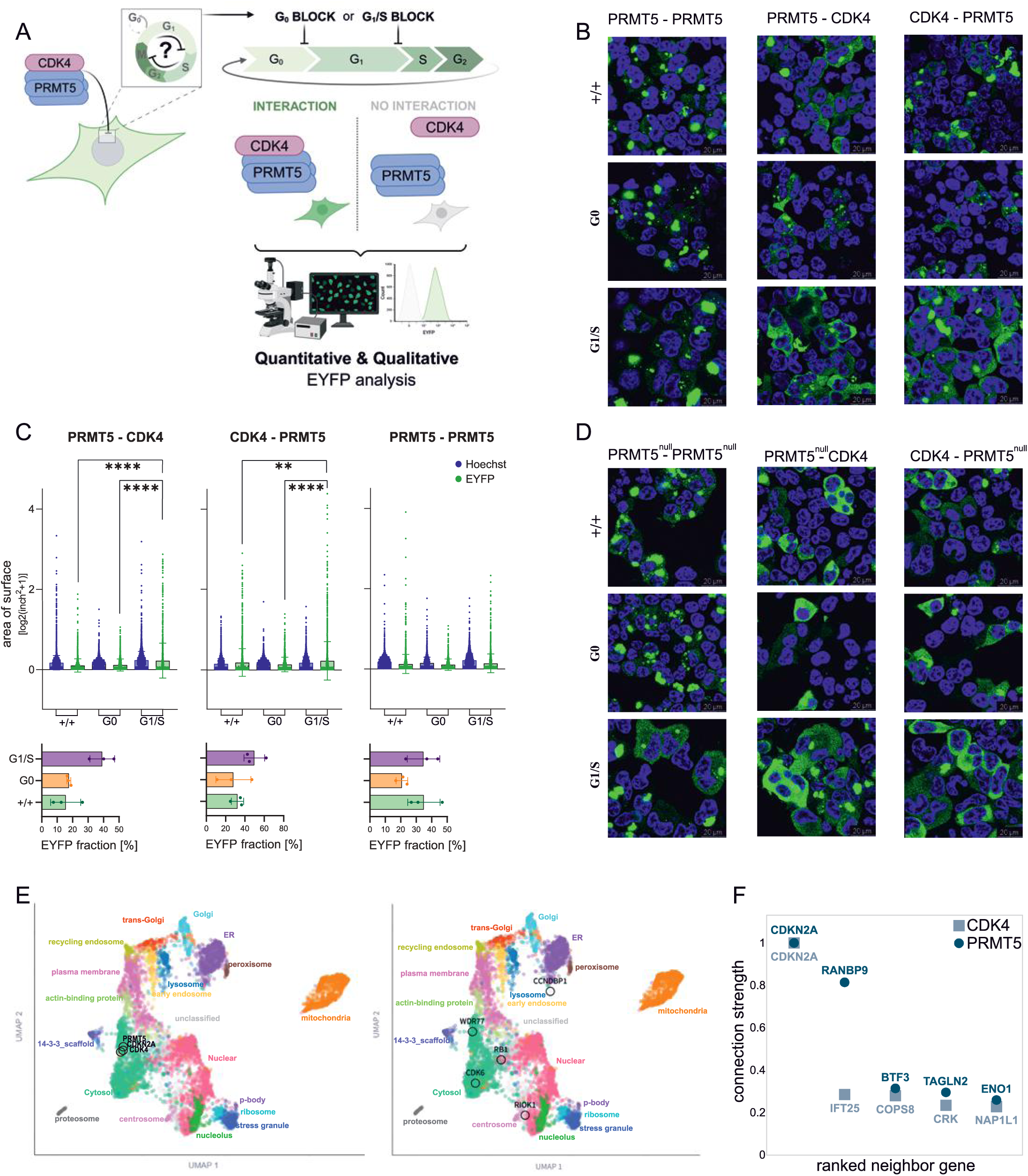
PRMT5 interacts with CDK4 in the cytosol in a cell cycle phase-dependent manner, independent of its methyltransferase activity. **(A)** Schematic overview of the workflow to investigate PRMT5–CDK4 interactions in a cell cycle–dependent context using a split EYFP complementation in living 293T cells, followed by confocal microscopy, quantitative image analysis, and flow cytometry (FACS). **(B)** Representative confocal microscopy images showing interaction complementation (EYFP signal) in living 293T cells expressing PRMT5^WT^ under the indicated cell cycle block conditions: +/+ (no block; DMEM with 10% FBS), G0 arrest via serum starvation (DMEM−/−), and G1/S arrest using a double thymidine block [2mM]. Nuclei were stained with Hoechst33342. Pairwise complementation was used to validate the interaction of selected protein pairs in living cells. **(C)** Quantification of one of three independent confocal microscopy experiments showing increased PRMT5–CDK4 interaction during G1/S arrest. EYFP signal area [log2(inch^2^+1)] and corresponding EYFP fractions [%] are presented in bar graphs. (Significance of upregulated EYFP-signal obtained via two-tailed Wilcoxon test with p-values adjusted for multiple testing using FDR correction * < 0.05, ** < 0.01, *** < 0.001, and **** < 0.0001). **(D)** Same analysis as in (**B**), performed with the catalytically deficient PRMT5^null^ mutant (G367A/R368A). **(E)** UMAP projection of protein subcellular localizations based on consensus graph-based annotations from the Organelle-IP database with proteins of interest highlighted in black circles. **(F)** Dot plot of the top five proteins with highest localization similarity scores (0–1) indicating the subcellular co-localization strength between PRMT5 and its neighbor proteins (dark blue circle), and CDK4 and its neighbor proteins (light blue square), respectively.

To corroborate these findings, we exploited the recently released high-resolution organelle immunocapture-MS data from 293T cells^31^ which established a map depicting the subcellular organization of more than 7,600 proteins. The two-dimensional UMAP, derived from graph-based data-driven annotation of protein localization, represents a spatial map of the HEK293T proteome with the arrangement of compartments reflecting known biological relationships. This map revealed a strong relationship between PRMT5 and CDK4 in the *bona fide* cytosol cluster (**Figure 3E, left**). In comparison, established complex partners or functionally related interactors of CDK4 (CCNDBP1, RB1, and CDK6) and of PRMT5 (WDR77 and RIOK1) were distant in the UMAP and, in some cases, annotated to different subcellular compartments (**Figure 3E, right**). Further, k-nearest-neighbor (k-NN) graph generated from the integrated subcellular proteomics dataset^31^, edge weights quantify pairwise subcellular localization similarity (connection strength, CS: 0 = no similarity, 1 = maximal similarity). Among the 1,836 proteins assigned to the cytosol, CDKN2A was identified as the top-scoring first-order neighbor of both PRMT5 (CS = 1) and CDK4 (CS = 1). This underscores their exceptionally similar cytosolic localization profile in this unbiased data set (**Figure 3F**). In more detail, the merged network graph comprising the first-degree neighbors derived from the organelle-IP data of PRMT5 and CDK4, revealed a subnetwork rich in cytosolic proteins and showed the overlap between the two interaction neighborhoods (**Figure S3D**). CDKN2A emerged as a central shared node, linking both proteins within a network primarily associated with cell cycle and mitotic regulation (**Figure S3E**).

### PRMT5 modulates CDK4 complex formation with CCND3 and CDKN2A independently of its methyltransferase activity

We next tested the impact of PRMT5 on CDK4 complex formation with CCND3 and CDKN2A/B. To investigate the potential interference of PRMT5, we modeled the CDKN2A–CDK4/6–CCND3 complex from the structural alignment of two complexes. We superimposed the CDK4–CCND3 structure (PDB: 7SJ3) and the CDK6–CDKN2A structure (PDB: 1BI7), generating a composite model with an RMSD of 2.769Å (**Figure 4A, top**) that shows the binding of CDKN2A and CCND3 on two opposite sides of the CDK4 N-lobe. Notably, the interaction surface on the N-lobe of CDK4 that engages with PRMT5 overlaps with the N-lobe binding region for both CCND3 and CDKN2A, however the majority of the residues form a bridge-like connection between the binding sites of two partners (**Figure 4A**). It has been shown, that binding of CDK4 to CCND1/3 drives nuclear translocation to phosphorylate nuclear substrates (e.g. RB1) and enable progression from G1 to S phase (*reviewed in^17, 54^*); however, if and how PRMT5 changes the binding of CDK4 and its partners and the complex dynamics is unknown. To experimentally test the hypothesis that PRMT5 modulates CDK4 complex formation (**Figure 4B**), we employed a sensitive NanoLuc-based LUMIER Co-IP reporter assay in a mammalian cell line ^33, 34^. At baseline (no PRMT), both CDK4–CCND3 and CDK4–CDKN2A interactions showed consistently strong binding signals as indicated through high luciferase activity after IP (log2FC of 4–6 relative to a non-binding control). Co-expression of CDKN2A/B or CCND3 with the respective CDK4–CCND3 or CDK4–CDKN2A complexes reduces complex formation, consistent with the reported mutual influence on binding of CCND3 and CDKN2A/B to CDK4 ^60, 61^. In our experiments, upon increasing PRMT5 levels through a dose-dependent titration of the PRMT5 plasmid (25–100ng) during transfection, the relative binding of CCND3 or CDKN2A to CDK4 progressively increased, reaching maximal binding signal at the highest PRMT5-amount (**Figures 4C**). Interestingly, this behavior was comparably observed for PRMT1 (**Figure S4**), in line with a report showing stabilized CDK4–CCND1 complex formation with PRMT1^28^. Since the assay employs whole-cell lysates, the observed enhancement of PRMT5 on CDK4 complex formation with CCND3 and CDKN2A, respectively, likely reflected a pool of nuclear and cytosolic CDK4 assemblies in different cell cycle phases and may also be indirect e.g. a result of competitive binding events. Nevertheless, the PRMT-dependent enhancement of the interaction dynamics occurred independently of their methyltransferase activity, as the PRMT5^null^ (**Figure 4D**) mutant recapitulated WT behavior. In addition to the PPI-complementation experiments highlighting that CDK4 and PRMT5 can form complexes in live cells, we used co-IP experiments to measure the impact of elevated PRMT5 expression on CDK4 interaction dynamics with its key regulators CCND3 and CDKN2A.

**Figure 4:**
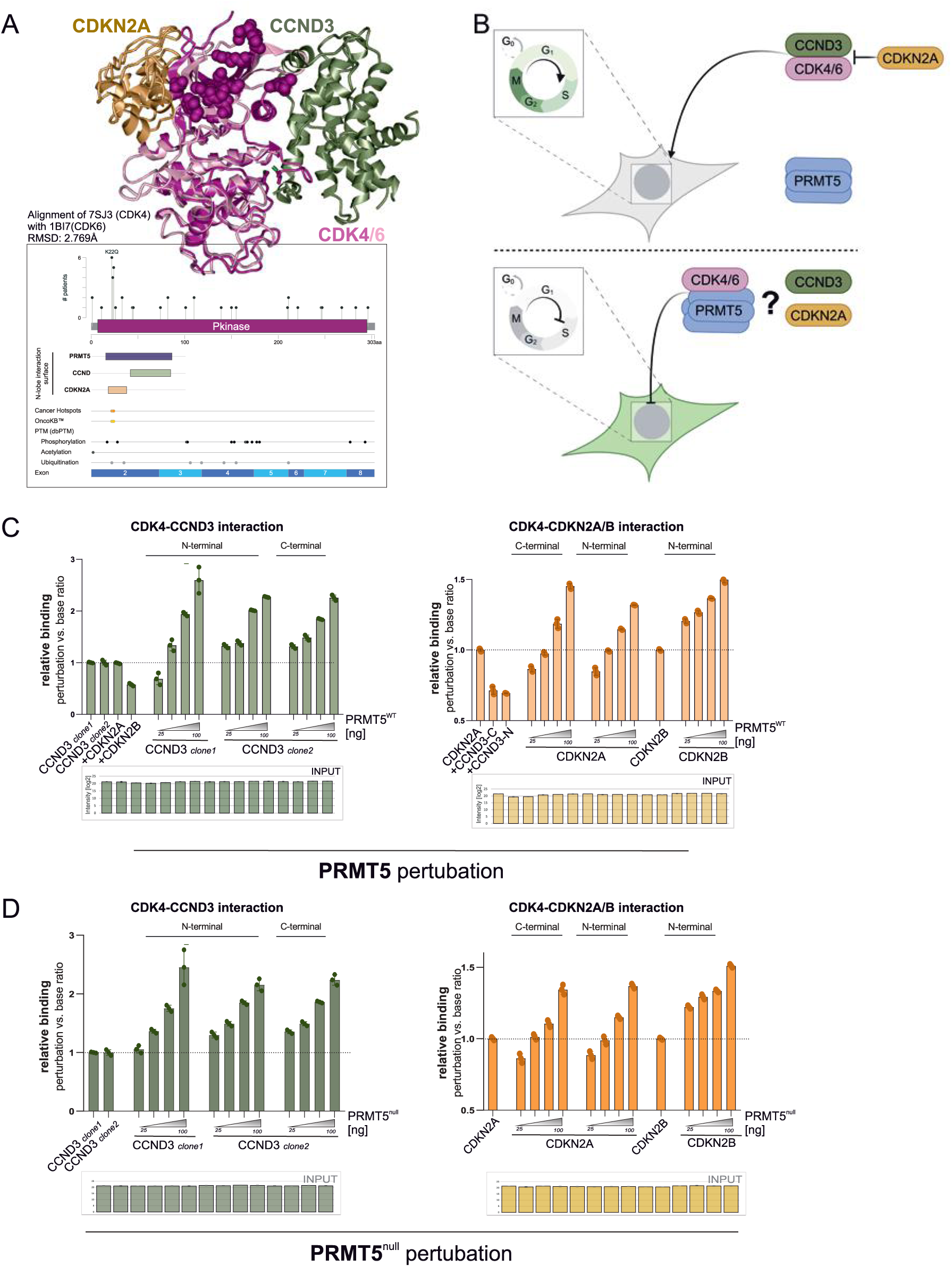
PRMT5 increases formation of the CDK4-CCND3 and CDK4-CDKN2A complexes independently of its methyltransferase activity. **(A)** Structural superimposition of CDK4–CCND3 (PDB: 7SJ3) and CDK6–CDKN2A (PDB: 1BI7) complexes (RMSD = 2.769Å) illustrates the central positioning of the predicted PRMT5 interaction interface on the N-lobe bridging the binding sites of CCND3 and CDKN2A. Cancer-relevant features, including functional domains, exons, hotspot mutations, and post-translational modification sites, were annotated using cBioPortal and visualized with mutation frequencies from TCGA data. **(B)** Schematic overview of the cell cycle highlighting the potential regulatory role of the PRMT5–CDK4 interaction in modulating cell cycle transitions via interference with activating CDK4–CCND or inhibitory CDK4–CDKN2A complexes. (**C**) Relative binding ratios from NanoLuc-based co-immunoprecipitation assays assessing the effect of PRMT5^WT^ perturbation on CDK4 interactions with CCND3 (green) and CDKN2A/B (orange). Data represent triplicates from three independent experiments, normalized to unperturbed controls. (**D**) Same analysis as (C), using the catalytically deficient PRMT5^null^ mutant (G367A/R368A).

### The phospho-proteomic response of PRMT5 overexpression mimics CDK4 inhibition

To better understand the functional impact of PRMT5 on kinase signaling, we performed quantitative phospho-proteomics. Using a doxycycline-inducible PRMT5-overexpressing cell line (Flp-In™ T-REx™ 293), we compared phospho-signaling profiles under four conditions: non-induced control, PRMT5 overexpression, CDK4/6i using Palbociclib (500 nM), and the combination of PRMT5 induction with CDK4 inhibition after 24h (**Figure 5A**).

**Figure 5:**
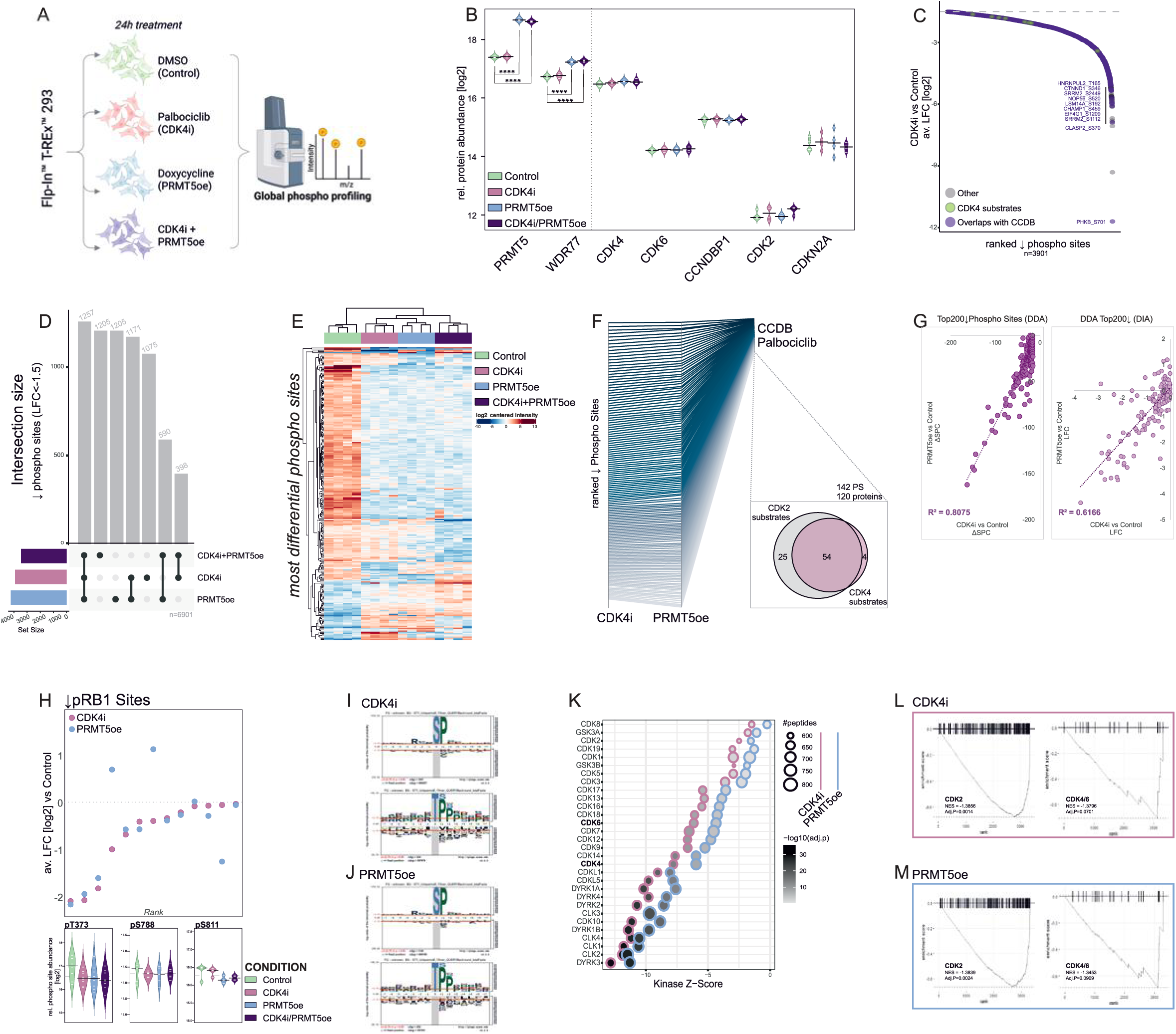
PRMT5 overexpression phenocopies/mimics CDK4 inhibition, with strongest phospho-suppression in RNA processing and cell cycle pathway. **(A)** Schematic overview of the phospho-proteomics workflow. Briefly, stable, inducible PRMT5-overexpressing Flp-In™ T-REx™ 293 cells were generated and treated for 24h under four conditions: 0.2% DMSO (control), 0.5µM Palbociclib (CDK4i), 2µg/mL doxycycline (PRMT5oe), or 0.5µM Palbociclib + 2µg/mL doxycycline (CDK4i + PRMT5oe). Following cell harvest, lysis, and SP3-based peptide purification, phospho-peptides were enriched using Zr/Fe-beads and subjected to phospho-proteomic profiling. **(B)** Normalized and transformed (log2) relative protein abundances of selected markers (C) in SP3-prepared input samples confirmed consistent protein levels across conditions, except for induced PRMT5 overexpression. (Significance obtained via two-tailed t-test with p-values adjusted for multiple testing using FDR correction * < 0.05, ** < 0.01, *** < 0.001, and **** < 0.0001). **(C)** Waterfall plot showing the most downregulated phospho-sites (log2FC < –1.5) in palbociclib-treated cells compared to DMSO-control, plotted as average log2FC relative to the control condition. Highlighted are overlaps with marked palbociclib-responsive sites derived from the Cell Cycle Database (CCDB) (violet) and annotated CDK4 substrates (green). **(D)** UpSet plot summarizing the most downregulated phospho-sites (log2FC < –1.5) compared to control across the three different conditions. **(E)** Heatmap representing the normalized log2-centered abundances of the most differentially regulated phospho-sites with Euclidean clustering across all treatment conditions. **(F)** Phospho-sites most downregulated (log2FC < −1.5) by CDK4i show substantial overlap with those affected by PRMT5oe and annotated Palbociclib-dependent downregulated sites from CCDB. Of 142 affected phospho-sites on 120 proteins, 79 (∼65%) are CDK2 substrates and 58 (∼48%) are CDK4 substrates, based on KEA3 set annotations with 54 common targets shown in the Venn diagram. **(G)** Scatterplots showing correlation between CDK4i and PRMT5oe for the 200 most depleted canonical phospho-sites based on ΔSPC from DDA data (site confidence >0.75, left), and matching sites from DIA data (site confidence >0.90, right), confirming consistent phospho-signatures across both datasets. **(H)** Waterfall plot of all downregulated phospho-sites on the RB1 protein CDK4i with the corresponding average log2FC upon CDK4i (pink) and PRMT5oe (blue) (top). Selected RB1 phospho-sites are well-established palbociclib-responsive phospho-sites shown individually upon the indicated treatments with their relative abundance level (bottom). **(I-J)** Probabilistic motif of the most downregulated phospho-sites (FG: log2FC < –1.5, BG: fasta-derived motives) for serine and threonine phospho-sites quantified in the overlap of DDA and DIA acquisition; CDK4i (n(S) = 1047; n(T) = 224) (I) and PRMT5oe (n(S) = 1144; n(T) = 252) (J). (**K**) Kinase enrichment analysis, utilizing the kinase activity and inference dashboard (KINAID), identifies downregulated CMGC kinases (z-score) based on phospho-sites (LFC < –1.5). Individual analyses for PRMT5oe (blue border) and CDK4i (magenta border) are shown, highlighting a remarkably similar kinase signature across both treatments. The number of mapped phospho-peptides is indicated by dot size, and the adjusted p-value by color intensity. (**L-M**) KSEA based on aggregated absolute substrate values derived from CDK4i (L) or PRMT5oe (M) for CDK2- and CDK4/6-substrates reveal highly similar enrichment profile and normalized enrichment score (NES) for both kinase sets.

In the quantitative proteome profile, we first confirmed the successful ∼ 4-fold induction of PRMT5 expression under doxycycline treatment. This was accompanied by an increased expression level of the endogenously co-regulated co-factor WDR77 (**Figure 5B**). Other significant protein expression chances were not observed, in particular protein levels of key cell cycle-related proteins (CDK4, CDK6, CDK2 etc.) were unaffected through any of the treatments after 24h (**Figure 5B, S5A**). Using Fe/Zr affinity enrichment of phospho-peptides we identified a total of 39,644 phospho-peptides from biological quadruplicates, corresponding to 32,149 sites quantified in at least one of the 4 conditions with high overlap between DDA and DIA acquisition mode (**Figure S5B**). 12,300 phospho-peptides were altered by more than 1.5 log2FC compared to control, whereby 2,467 to 2,808 increased and 3,450 to 4,223 phospho-peptides were downregulated per condition.

First, we validated the effect of Palbociclib-mediated inhibition on the phospho-proteome. As a reference we utilized phospho-proteomic data from a recent study that employed Palbociclib treatment and phospho-proteomics and established a set of 401 cell cycle response proteins with 6,528 regulated phospho-sites (Cell Cycle Database CCDB^62^). We cross-referenced the phospho-sites that were downregulated in our dataset (log2FC < –1.5) with those annotated as Palbociclib-responsive in the CCDB. Of the 3,901 phospho-sites reaching our significance threshold in the CDK4i condition, 2,476 (>60%) overlapped with annotated Palbociclib-responsive sites, supporting that our observed phospho-site changes robustly reflect a CDK4/6 inhibitory effect (**Figure 5C**). Notably, 2,428 phospho-sites were most downregulated (log2FC < –1.5) in both CDK4i and PRMT5oe (**Figure 5D**). The global phospho-proteomic profiles, when clustered and displayed as heatmap, revealed strikingly similar patterns of deregulated phospho-sites upon CDK4i and PRMT5oe (**Figure 5E**). Thus, elevated PRMT5 expression and CDK4/6 inhibition induce strikingly similar phospho-proteomic signatures, suggesting functional convergence on the same signaling network. 142 phospho-sites that were downregulated in the reference CCDB-Palbociclib treatment overlapped with sites that were downregulated in both conditions, CDK4i and PRMT5oe (**Figure S5C**). Log2FC rank-based comparison across these conditions revealed a qualitative and quantitative similarity of the phosphorylation profiles with the CCDB-Palbociclib downregulated reference sites (**Figure 5F**). These 142 downregulated phospho-sites map to 120 proteins, of which 79 (∼65%) are annotated as CDK2 substrates and 58 (∼48%) as CDK4 substrates^63^ with 54 proteins that are common targets of both kinases (**Figure 5F**, Venn diagram). Moreover, quantitative assessment of the top 200 downregulated phospho-sites in the pharmacological CDK4i and interaction-dependent PRMT5oe experiment by either *i)* comparison of spectral counts (SPC) from our DDA-derived data or *ii)* by correlation of log2FC intensity values from DIA-derived data demonstrated a strong correlation between CDK4i and PRMT5oe conditions (R^2^ = 0.8079 and R^2^ = 0.6166 respectively, **Figure 5G**). RB1 is the most prominent substrate of CDK4, and RB1 phosphorylation marks the R checkpoint into G1/S of the cell cycle. This makes RB1 an ideal example to highlight the phospho-response mimicry between the treatments. CDK4i and PRMT5oe similarly downregulated a range of RB1 phospho-sites (**Figure 5H, top**), including target sites that are known to respond to Palbociclib-CDK4/6i, like T373, S788, and S807/S811 (**Figure 5H, bottom**). The sites are coupled to indirect CDK2 suppression^64^. Loss of S807/S811 phosphorylation leads to RB1 reactivation and G1 arrest^65, 66^, reduced T373 phosphorylation enhances suppression of E2F-driven proliferation^65^, and dephosphorylation at S788 disrupts E2F–RB1 complex dissociation, further reinforcing cell cycle arrest^67^. More globally, we obtained almost identical pS- and pT-centered linear phospho-site motifs for CDK4i and PRMT5oe downregulated phospho-sites (**Figure 5I-J, S5D-E**). The motifs highlighted a strong preference for proline at the +1 position next to the phospho-site, indicating a kinase substrate preference for proline directed kinases such as CDKs^68^.

Additionally, equal kinase motif scores were derived across kinase families with CMGC kinases as top downregulated kinase family (**Figure S5F**). When analyzing the kinase signatures of the most downregulated phospho-sites (log2FC < -1.5) with the recently published KINAID (Kinase Activity and Inference Dashboard)^69^,we observed a striking similarity between CDK4i and PRMT5oe condition with respect to phospho-signature enrichment for many members of the CMGC kinase family, including CDK4 and CDK6 (**Figure 5K**). Specifically, among the 10 most depleted kinase substrate enrichment (KSEA) signature sets (**Figure S5G**), CDK4/6 and CDK2 phospho-site set enrichment profiles for PRMT5oe and CDK4i were highly similar (**Figure 5L-M**). Together, the phospho-proteomics data provides a deep functional signaling readout underscoring that PRMT5oe mimics the cellular effects of CDK4i. This is in agreement with the findings demonstrating that enhanced PRMT5–CDK4 interaction in the cytosol negatively regulates CDK4 function and leads to G1 accumulation within the cell cycle.

## DISCUSSION

PRMT–kinase interactions with their broad capacities to regulate cellular signaling processes are particularly relevant molecular connections in cancers. To date, only a few PRMT–kinase interactions have been identified and characterized^26–30^. By mapping PRMT–kinase interactions in a systematic Y2H matrix approach, we uncovered a binary interaction network between 4 PRMTs and 20 protein kinases, many from the CMGC kinase family. The set of PRMT5 interacting kinases, revealed a functional kinase signature associated with the cell cycle. (**Figure 1**). Co-expression (**Figure 2A**) and interactome overlap analyses (**Figure 2B-D**) prioritized CDK4 as a top PRMT5 interacting candidate for further functional characterization. We obtained high scoring structural models predicting direct interaction of CDK4 with PRMT5 as part of a hetero-multimeric complex assembly (**Figure 2E–G**). We then studied the PRMT5 binding to CDK4 in live-cells and observed that the CDK4-PRMT5 complex localized in the cytosol. Applying cell cycle blocks, the interaction was increased in cells during the G1/S phase **(Figure 3)**. Notably, all experiments mechanistically point towards a methyltransferase activity independent function of the complex. The binding of CDK4 regulatory proteins is key to its activity^17, 54^. The active CDK4–CCND3 complex translocates to the nucleus^70, 71^, while association of the inhibitory CDK4–CDKN2A complex is primarily observed in the cytosol^71, 72^. The two regulatory partners mutually destabilize the other’s interaction^73, 54^. Importantly, entry of CDK4 to the nucleus is a critical step in promoting the cell cycle during G1/S transition. We hypothesize that the PRMT5 interaction retains CDK4 in the cytosol, delaying its nuclear function of phosphorylating RB1 and thus high levels of PRMT5 may contribute to the CDK4/RB1 checkpoint control in an inhibitory manner. This may actually be accentuated in our experimental setup, as EYFP interaction complementation can be induced through cellular stimuli such as the cell cycle phase, however upon interaction assisted EYFP folding dynamic properties of the system are lost^55^. While this mechanism aligns with previous findings in hepatocellular carcinoma^74^, it is important to note that this earlier report suggests a competitive interaction between PRMT5 and CDKN2A for CDK4 binding to regulate G1/S progression. Considering other apparently conflicting data that *(i)* show that nuclear PRMT5 can impact the CCND–CDK4 complex^70, 75^ and *(ii)* that the localization of PRMT5 and CDKN2A are inversely correlated^71^, we turned to recent high-resolution MS-based data from in organelle-specific IPs. In the Hein *et al.* subcellular localization map^31^ the cytosolic co-occurrence of PRMT5, CDK4 and CDKN2A is strikingly clear (**Figure 3E–F, S3D**). Furthermore, in luciferase-based co-IP assays we observed a PRMT5 dose-dependent enhancement of both the CCND3–CDK4 and CDKN2A–CDK4 complex formation (**Figure 4C–D**). The structural interface model, where PRMT5 contact residues in the CDK4 N-lobe bridge between the two binding regions is compatible with our observations in the binding assays. However, we do not exclude that the effects are indirect, related to competitive binding events possibly impacting the interplay with additional endogenous binding partners. These findings corroborate a regulatory potential of PRMT5 in allosterically modulating CDK4 interaction and localization dynamics by fine-tuning likely nuclear activation through increase in cytosolic interaction. This hypothesis was scrutinized through phospho-proteomic profiling, providing a data rich functional readout. We revealed that PRMT5 overexpression phenocopies the effects of pharmacological CDK4i (Palbociclib). The striking resemblance of phospho-site downregulation of CDK4i to PRMT5oe was demonstrated through clustering of global phospho-profiles, quantitative comparison with a known cell cycle responsive set of phospho-sites and through kinase substrate profiling (**Figure 5**).

In recent reports, PRMT5 inhibition led to alterations of cellular cell cycle profiles. siPRMT5 treatment blocks G1-to-S cell cycle transition in an RB-independent manner^76^. When monitoring cell cycle progression in the presence of the PRMT5 inhibitor GSK3326595, an interesting heterogeneity was observed across cancer cell lines: while a subset exhibited an increased proportion of cells in the G1 phase, others showed no apparent response, and yet another group displayed a decrease in the G1 phase population upon treatment^77^. Further, in T helper 1 (mTh1) cells, the PRMT5 inhibitor, HLCL65, arrested activation-induced T cell proliferation at the G1 stage of the cell cycle^78^. These reports highlight the transcriptional and also the splicing effects of PRMT5 on the p53/MDM4 or FUS/Pol II axis that may trigger cell cycle changes. PRMT5 localization itself has also been linked to cell-cycle control, where cytosolic PRMT5 drives proliferation and oncogenic transformation, while nuclear PRMT5 accompanies cell-cycle arrest, differentiation, and growth inhibition^78–80^. These effects may not involve CDK4, in the nucleus CCND1–CDK4 enhances PRMT5 activity by phosphorylating its co-factor WDR77^70^, which leads to recruiting PRMT5 to the chromatin via the adaptor COPR5 and CCNE1 transcription repression^75^. Moreover, PRMT5 inversely correlates with CDKN2A in RNA-seq data of oropharyngeal squamous cell carcinoma patients, with high cytosolic PRMT5 linked to elevated nuclear and reduced cytosolic CDKN2A^71^. While these observations are likely dependent on the specific cell context, together this underscores PRMT5’s functional connection with CDK4/6 signaling. In cancers with *CDKN2A/MTAP* deletions, both PRMT5 as well as CDK4/6 inhibition play a role in first line therapies. Resistance to CDK4/6i often involves failure to suppress PRMT5 activity, allowing escape via alternative cell cycle drivers, as PRMT5 activity functionally compensates for CDK4/6 loss^24, 76, 77^. Therefore, specifically in CDK4/6i–resistant tumors, inducing or sustaining PRMT5 hyperactivation, as alternative to PRMT5i^24, 76^, may lead to selective vulnerabilities.

In conclusion, our findings demonstrate direct physical interaction between PRMT5 and CDK4 in the cytosol and provide evidence that PRMT5 functions as a negative regulator of CDK4 activity. PRMT5 overexpression mimics CDK4 inhibition, disrupting both cellular signaling and cell cycle progression in a highly similar manner. These insights establish PRMT5 as a mediator and modulator of CDK4-driven cellular oncogenic programs, offering a novel rationale for exploiting PRMT5 expression and activity in cancers therapeutic strategies.

## Methods

### Cloning procedure and construct generation

All open reading frames (ORFs) used in this study were either obtained in gateway-compatible entry vectors or PCR-amplified from cDNA templates and subsequently cloned into entry vectors via BP recombination (Invitrogen). ORFs were transferred into destination vectors using LR recombination reactions, following the manufacturer’s standard Gateway protocol (Invitrogen). For yeast two-hybrid (Y2H) assays, ORFs were cloned into LexA-DB-based bait vectors (pBTM116-DM and pBTMcC24-DM) and GAL4-AD-based prey vectors (pACT4-DM and pCBDU-JW)^43^. For mammalian NanoLuc-based interaction assays, bait constructs were cloned into a Protein A fusion vector (pcDNA3.1PA-D57)^34^ and prey constructs into a NanoLuc-fusion vector (pNL-DM, courtesy of Taipale lab (Toronto)). For EYFP-Venus complementation assays, ORFs were cloned into split EYFP vectors^57^ containing either N-terminal or C-terminal Venus fragments fused to candidate proteins. All constructs were initially verified by restriction digest and select constructs, including all point mutants, were further confirmed by Sanger sequencing.

| CloneID | UniProtKB | Symbol | NCBI GeneID |
| --- | --- | --- | --- |
| GC-Y3476-CF | Q86X55 | CARM1 | 10498 |
| GC-Y3476 | Q86X55 | CARM1 | 10498 |
| RZPDo839F06135 | P30281 | CCND3 | 896 |
| CCSB_6197 | P30281 | CCND3 | 896 |
| CCSB_4019 | P11802 | CDK4 | 1019 |
| CCSB_13668 | Q00534 | CDK6 | 1021 |
| CCSB_7698 | P42772 | CDKN2B | 1030 |
| CCSB_56119 | Q99873 | PRMT1 | 3276 |
| RZPDo834A082D | Q99873 | PRMT1 | 3276 |
| RZPDo839F0381 | O14744 | PRMT5 | 10419 |
| RZPDo834G0114D | O14744 | PRMT5 | 10419 |
| MWM1025 | Q96LA8 | PRMT6 | 55170 |
| CCSB_55527 | Q96LA8 | PRMT6 | 55170 |
| RZPDo839E12147 | Q9NR22 | PRMT8 | 56341 |

### Site directed mutagenesis

PCR amplification of PRMT1 and PRMT5 gene fragments was performed using Phusion High-Fidelity DNA Polymerase (NEB, M0530S). Each PCR reaction contained 1× Phusion HF buffer, 200µM dNTPs (NEB, N0447S), 0.5µM each of forward and reverse primers, 3% DMSO, 0.02U/µL Phusion DNA Polymerase, ∼0.35ng/µL template DNA, and nuclease-free water to final volume. The thermocycling protocol included initial denaturation at 98 °C for 1min, followed by 30 cycles of 98°C for 30s, 67°C for 30s, 72°C for 2min, and a final extension at 72°C for 10 min. PCR products were digested with DpnI (NEB, R0176S) by adding 2µL DpnI mix (0.2µL DpnI, 1.2µL CutSmart Buffer, 0.6µL H₂O) to 10µL PCR product, incubating at 37°C for 3.5h, and inactivating at 80°C for 20min. Following chemical transformation in *E.Coli*, three colonies were screened via colony PCR and verified by Sanger sequencing. Final inserts were subcloned via LR recombination into destination vectors.

PRMT5_G367A/R368A_fwd:

5’ GTGCTGGGAGCAGCCGCAGGACCCCTGGTGAACGCTTCCCT 3’,

PRMT5_G367A/R368A_rev:

5’ CACCAGGGGTCCTGCGGCTGCTCCCAGCACCATCAGTACCTGGAA 3’

PRMT1_W215L_fwd:

5’ GACTACAAGATCCACCTCTGGGAGAACGTGTATGGCTTCGACATGTCT 3’,

PRMT1_W215L_rev:

5’ ATACACGTTCTCCCAGAGGTGGATCTTGTAGTCTTTGTACTGCC 3’

### Yeast Strains, growth, and transformation

Yeast strains L40ccU2 (MATa) and L40ccα (MATα) were used for bait and prey expression, respectively^45, 34^. Cultures were maintained on YPDA medium (1% yeast extract, 2% peptone, 2% glucose, 0.004% adenine) or on nitrogen base (NB) medium supplemented with 2% glucose, 0.004% adenine, and 0.002% uracil. Selective supplements were added according to experimental selection needs: bait selection (Leu, His), prey selection (Trp, His), mating controls (His), and auto-activity tests (Leu/ Trp).

Yeast transformation was performed following an adapted lithium acetate/single-stranded carrier DNA/PEG protocol. Briefly, overnight cultures grown in YPDA at 30°C were diluted to OD_600_ 0.1-0.15 and expanded to mid-log phase. Cells were harvested, washed, and resuspended in transformation buffer (0.1M lithium acetate, 5mM Tris-HCl pH 8.0, 0.5mM EDTA, 1M sorbitol). Plasmid DNA (350-500ng) and salmon sperm DNA (25µg) were added to each transformation. Following incubation with PEG-based solution, DMSO addition, and heat shock at 42°C, transformed cells were plated in triplicate and treated separately as biological replicates.

### High-Throughput Yeast Two-Hybrid Interaction Screening

Binary interactions were systematically mapped using a high-throughput Y2H matrix approach in a 384-colony format, as previously described^45, 34^. Active kinase libraries^39, 40^ were transformed into yeast strains and subjected to automated robotic mating using a K8 arrayer (Kbiosystems Limited, UK). Using robotics support, bait strains cultured in liquid medium were mixed with prey matrices grown on selective media and mated on YPDA plates. Diploids were selected on SD medium (lacking leucine, tryptophan, adenine, uracil, and histidine) solidified with 2% agar. Colony growth was monitored daily over seven days at 30°C, and interaction colony size was scored by visual inspection. Only interactions confirmed across biological replicates were included in the analysis. Auto-activating bait constructs, identified by growth on -His media without mating, were excluded from the final analysis. Raw data and processed scoring tables can be found in the supplemental section (**Suppl. Table 1-2**).

### Computational Analysis

Co-expression correlation analysis and kinase enrichment analysis were performed using the ARCHS4 gene expression datasets via Enrichr^49^. The top 100 genes co-expressed with the target gene were used as the query set (**Suppl. Table 3**). Statistical enrichment was evaluated based on adjusted p-values obtained from Enrichr.

The subcellular localization of PRMT5 and CDK4 was analyzed computationally using publicly available organelle proteomics data^31^. Enrichment profiles from native organelle IP-MS in HA-tagged HEK293T cells were accessed via organelles.czbiohub.org/ Subcellular_UMAP (*accessed 25-10-26*) and used to assign localization based on graph-based clustering (**Suppl. Table 4**).

To validate the effectiveness of CDK4 inhibition by Palbociclib, a curated set of palbociclib-responsive phospho-sites was obtained from the Cell Cycle Database^62^. Only sites showing regulation after 24h of Palbociclib treatment and were annotated upon peak average as “Palbo”, at the timepoint immediately following release (t = 0), were selected to match the conditions used in this study’s phospho-proteomic analysis (**Suppl. Table 6, 8**). These phospho-sites served as functional readouts of CDK4 activity and were used to evaluate the cellular response to treatment.

### AlphaFold-Multimer Structure Prediction

Protein-protein complex structures were generated using AlphaFold-Multimer^32^, conducted via the ColabFold interface^81^ and on a local installation provided by collaborators at Aarhus University. FASTA sequences of target proteins were used as inputs. Predicted structures were analyzed based on pLDDT score. Structures were visualized using iCn3D^82^, with interface confidence assessed per model confidence metrics. PDB files are available in the Supplement.

### Confocal microscopy and quantification of EYFP-Venus complementation

HEK293T cells (1×10^5^/well) were seeded in poly-D-lysine–coated 24-well black glass-bottom plates (ibidi, µ-Plate Black) and subjected to a sequential cell cycle synchronization protocol. To this end, cells were either serum-starved for 24h in DMEM without FBS to induce G0 arrest, treated with 2mM thymidine (Sigma-Aldrich, T9250) for 20–24h to block cells at the G1/S boundary, or left untreated in complete medium (DMEM + 10% FBS) as a control. After 24h and before transfection, cells were released from the respective blocks. EYFP-Venus split complementation constructs were transfected using 250ng per plasmid per well through the PolyJet transfection reagent mix (SignaGen, SL100688) at a 1:3 (w:v) DNA:PolyJet ratio. 6h post-transfection, the medium was replaced with fresh synchronization medium corresponding to the original condition (serum-free, thymidine, or complete) to initiate a second block for an additional 18–20h.

For nuclear staining, cells were incubated with Hoechst33342 (1µg/mL; Thermo Fisher, H3570) in the respective blocking medium. Confocal imaging was performed using a Leica Stellaris 5 system equipped with a 63x/1.4 NA oil immersion objective. Fluorophores were excited at 405nm (Hoechst) and 514nm (Venus EYFP), and images were acquired using LAS X software under identical settings across all samples. EYFP complementation signals and subcellular distribution were quantified using a semi-automated ImageJ macro. Image channels were split, and fluorescence areas were measured based on surface coverage. Nuclei were segmented via Hoechst staining, and PPI was assessed by calculating the EYFP-positive area relative to the nuclear area. A log₂-transformed Hoechst-to-EYFP ratio (log₂-transformed (inch2+1)) was computed from the measured fluorescence areas. For each complementation condition, a minimum of three representative fields encompassing at least 600 nuclei or objects were analyzed.

### NanoLuc-based Co-immunoprecipitation Assay

LUMITRAC 96-well plates were coated by incubating each well with sheep γ-globulin (1:1000; Jackson ImmunoResearch, 013-000-002) for 24h at 4°C, followed by blocking with 2% BSA for an additional 24h at 4°C. Subsequently, wells were incubated with AffiniPure Rabbit Anti-Sheep IgG (H+L) (1:750; Jackson ImmunoResearch, 313-005-003) for 24h at 4°C to facilitate Protein A-mediated capture. HEK293T cells (ATCC®, CRL-3216™) were cultured in DMEM supplemented with 10% FBS and 1% PenStrep at 37°C in a humidified atmosphere with 5% CO₂. Cells (2×10⁴/well) were seeded in poly-D-lysine–coated 96-well plates (0.05mg/mL, Sigma-Aldrich, P7405) and co-transfected in triplicates using PolyJet (Signa Gen, SL100688) at a 1:3 (w:v) DNA:PolyJet ratio. Each well received a total of 150ng DNA, consisting of 25ng Nanoluc, 25ng PA-construct, and 0-100ng perturbation plasmid. The total was adjusted to 150 ng using an empty vector to ensure equal DNA concentration across all wells. 48h post-transfection, cells were lysed with SDS-free Hengs buffer (50mM HEPES-KOH, pH 7.9; 150mM NaCl; 20mM Na₂MoO₄; 2mM EDTA; 5% glycerol; 0.5% Triton X-100) supplemented with ProtI and PhosI for 3h at 4 °C, shaking. Cleared lysates (80µL) were added to the prepared IgG-coated plates and incubated for 2h at 4 °C upon gentle shaking to allow capture of PA-tagged bait and associated NanoLuc-fused prey. Wells were washed three times with 100µl ice-cold inhibitor-free lysis buffer, followed by the addition of 40µL PBS and 40µL Nano-Glo® Luciferase Assay Reagent (Promega, N1130). Luminescence was recorded at the luminescence emission of 460nm sequentially at room temperature after 0.5-2h using a DTX880 microplate reader (Beckman Coulter). All measurements were performed in triplicate transfections. Mean log₂-transformed raw luminescence signals and standard deviations were calculated for each DNA construct pair. Values were normalized to background signals from non-binding PA-fusion controls or to unperturbed PA-fusion reference levels. Interaction signals were considered significant if the normalized ratio exceeded 2-fold over background and exhibited a z-score > 2.

### Generation of Flp-In™ T-REx™- inducible PRMT5 overexpressing cell lines

Stable, inducible Flp-In™ T-REx™ 293 cell lines were generated using an adapted workflow of the established protocol from the manufacturer Invitrogen. Briefly, The PRMT5-inducible Flp-In™ T-REx™ system was generated by co-transfecting cells with a 1:9 ratio of pcDNA5/FRT/TO and pOG44 (0.48µg/mL total DNA) using PolyJet™. After 5h, medium was replaced with DMEM + 15µg/mL blasticidin. Selection began 24h later with 15µg/mL blasticidin and 50µg/mL hygromycin B. Polyclonal populations were expanded and validated by immunoblotting and MS.

### SP3 sample preparation for phospho-proteomics

Flp-In™ T-REx™ 293 cells with inducible PRMT5 overexpression were seeded and treated the following day with DMSO (vehicle control), 0.5µM Palbociclib (MedChemExpress, 571190-30-2), 2µg/mL doxycycline (Fisher Scientific, 10224633), or a combination of Palbociclib and doxycycline for 24h. All treatment conditions were adjusted to a final DMSO concentration of 0.21%. Following harvest in ice cold 1xPBS, cell pellets were lysed in 2.5% SDS, 50mM AmBic, +ProtI +PhosI. Post determination using BCA following the manufacturer’s instructions, protein concentration was adjusted to 1mg/ml using 100mM AmBic, and lysates were pre-cleared at 10 °C for ≥15 min at 14,000xg. Proteins were reduced by 5mM TCEP (Merck, 646547-10X1M) and alkylated with 10mM chloroacetamide (Pierce, A39270), each for 10min at 37°C in the dark. The SP3 procedure was adapted from the previously reported method^83^. Briefly, SP3 cleanup was performed by adding Sera-Mag SpeedBeads (Merck, EM1 100/40) at a 1:5 (w:w) bead-to-protein ratio, followed by acidification to 0.4% FA and addition of ACN to a final concentration of 50%. After 10min at RT, beads were magnetically separated and washed twice with 70% EtOH and once with 100% ACN. Beads were air-dried (<5min) and digested with trypsin (Fisher Scientific 13454189) in 100mM AmBic at a 1:100 enzyme-to-protein ratio for 2–4 h at 37°C shaking, followed by overnight digestion with the same ratio. Peptides were generated and released from the SP3 beads via on-bead tryptic digestion in 100 mM ammonium bicarbonate. Following digestion, samples were acidified to 2% TFA and lyophilized for ≥ 12h. An aliquot (∼10%) was retained as an unenriched input control.

### Phospho-peptide enrichment using Zr-IMAC and Fe-NTA magnetic beads

The method was developed by following established protocols^84, 85^. Dried peptides were resuspended in 50µL freshly prepared binding buffer (80% ACN, 5% TFA, 0.5% glycolic acid). For every 100µg peptide, 10µL Zr-IMAC MagReSyn HP beads (PELOBiotech, MR ZHP005) and 2.5µL Fe-NTA MagBeads (PureCube, 31501-Fe) were pre-washed three times in binding buffer and resuspended in 20µL binding buffer. Samples were sonicated at 4°C for 10 min (30s ON/OFF) followed by pre-clearance at 4°C and incubated with the mixed beads for 20–30min at RT on a rotating wheel. Beads were washed sequentially with 200µL each of binding buffer, wash buffer I (80% ACN, 1% TFA), and wash buffer II (10% ACN, 0.2% TFA). Phosphopeptides were eluted twice using 150µL freshly prepared 1% ammonium hydroxide, combined, neutralized with 100µL 10% FA, and lyophilized for ≥24h followed by SpeedVac at 37°C for 1h. Lyophilized peptides were resuspended in 0.1% FA. Input samples were quantified and diluted to 200ng/µL in 0.1%FA for LC-MS/MS injection.

### Mass spectrometry

Samples were analyzed on a timsTOF Pro ion mobility mass spectrometer (with PASEF® technology, Bruker Daltonics) in line with UltiMate 3000 RSLCnano UHPLC system (Thermo Scientific). Peptides were separated on a reversed-phase C18 Aurora column (25cm × 75µm) with an integrated Captive Spray Emitter (IonOpticks). Mobile phases A 0.1% FA in water and B 0.1% FA in ACN (Fisher Scientific, 10799704) with a flow rate of 300nL/min, respectively. Fraction B was linearly increased from 2% to 25% in the 90min gradient, increased to 40% for 10min, and a further increase to 80% for 10min, followed by re-equilibration. The spectra were recorded in DDA and DiaPASEF and analyzed with MSfragger^86^. DDA data were used for spectral library generation using MSfragger (v22.0) in library-free mode against UniProt-reviewed human proteome (*download: 20.01.2025*). Raw DIA data were processed using the generated library in DIA-NN 2.1.0.^87^. DDA and DIA Perseus-imputed data were normalized (Variance Stabilizing Normalization) and log2-transformed using FragPipe^88^. Downstream statistical analysis and visualization were performed with Access, Cytoscape, Enrichr, FragPipe-Analyst, and R.

### Kinase enrichment

To perform enrichment analysis the previously determined phosphorylation profile of the PRMT5oe condition and CDK4i were first simplified (**Suppl. Table 7**). For each protein, the phosphorylation changes across all quantified peptides were consolidated by calculating the absolute sum of the log2FC values. These proteins were then ranked in decreasing order based on their absolute sum of phosphorylation changes. The complete sets of known substrates for all kinases belonging to the CMGC and CAMK families were systematically extracted from the PhosphoSitePlus (version 210217) was performed to identify kinases whose substrates were significantly enriched among the downregulated phosphorylation sites. This analysis was conducted using the fgsea Gene Set Enrichment Analysis (GSEA) package^89^ in R. Only kinase-substrate sets comprising more than 40 substrates were included in the enrichment analysis to ensure statistical robustness. Visualization of the resulting enrichment data was executed in R using the ggplot2 package.

### KINAID and Kinase motifs

Differentially regulated phospho-sites identified from quantitative phosphoproteomic analysis were used for kinase activity and motif inference. Phospho-sites with a log2FC < –1.5 (**Suppl. Table 7**) were selected as the most strongly downregulated sites and analyzed using KINAID ^69^ to predict kinase activity patterns and phospho-signature enrichment based on curated kinase–substrate interaction databases. The resulting profiles revealed strong enrichment across multiple members of the CMGC kinase family. To identify consensus phosphorylation motifs, sequences surrounding each modified residue (±6 amino acids) were compared against curated kinase substrate motifs from PhosphoSitePlus® (https://www.phosphosite.org/, *accessed 06/2025*). Further, probability motif logo analysis was performed using pLogo ^90^, with all DDA-identified phospho-sites as the background dataset. Separate analyses were conducted for serine (pS) and threonine (pT) sites to visualize residue-specific sequence preferences and statistically enriched amino acid patterns around phosphorylation sites.

## Supporting information

pdb_01

pdb_02

## Data availability

The predicted AlphaFold structures can be found together with the Suppl. Tables in the supplementary section indicated as PRMT5–WDR77–CDK4_deca.pdb and PRMT5–WDR77–CDK4_dodeca.pdb, respectively. The mass spectrometry data generated in this study have been deposited via the ProteomeXchange PRIDE partner repository ^91^ with the dataset identifier PXD065510.

## Figure Legends

**Figure S1:**
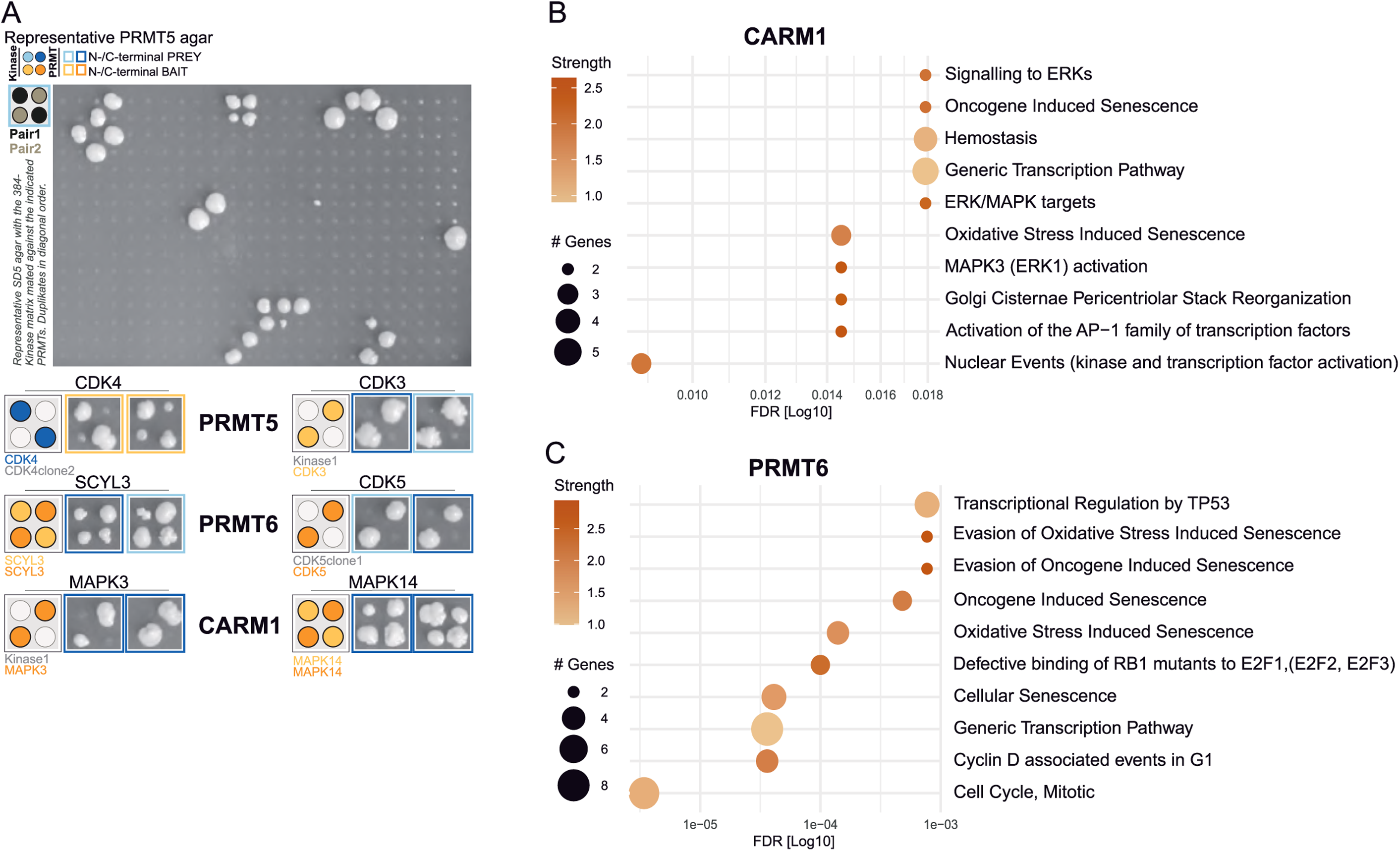
Establishment of a comprehensive PRMT–Kinase Network indicates the importance of PRMTs in cell cycle-associated processes. (**A**) Representative selective agar (SD) plates (lacking Leu, Trp, Ade, Ura, and His) from yeast two-hybrid (Y2H) screening of a 384-kinase matrix against the indicated PRMT constructs after seven days of incubation. Interaction partners were identified by duplicate colony growth. (**B-C**) Functional enrichment analyses of uniquely identified PPIs for CARM1 (B) and PRMT6 (C), showing the top 10 enriched biological pathways based on Reactome annotations, ranked by false discovery rate (FDR) using String-DB analysis.

**Figure S2:**
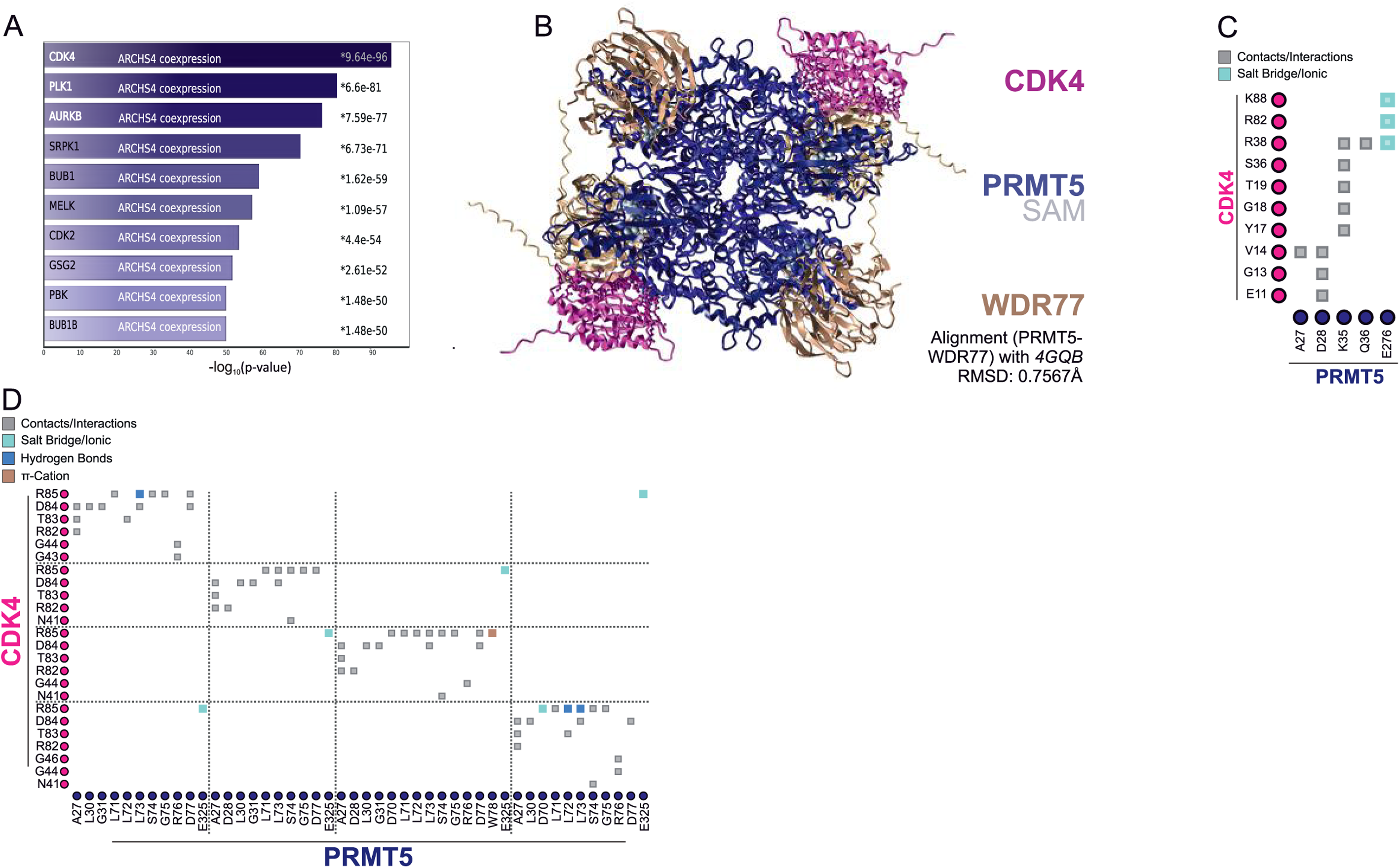
Data integration reveals the importance of a hetero-multimeric PRMT5-CDK4 complex in cell cycle regulation. (**A**) Kinase gene set co-expression analysis based on the top 100 genes co-expressed with PRMT5, identifying the most significantly correlated kinase gene sets using ARCHS4 RNA-seq data derived from Enrichr. (**B**) Structural alignment of the AlphaFold-Multimer–predicted PRMT5–WDR77–CDK4 complex (generated in duplicate) with the experimentally resolved hetero-octameric methylosome core complex (PDB: 4GQB), comprising four PRMT5 and four WDR77 subunits bound to SAM. The alignment yielded a root-mean-square deviation (RMSD) of 0.7567Å, indicating high structural similarity. (**C**) Interaction map detailing amino acid contacts and interaction types between a single PRMT5–CDK4–WDR77 unit from the predicted complex in (**B**). (**D**) Comprehensive interaction map showing contact residues and interaction types among all PRMT5 and CDK4 molecules within the predicted hetero-dodecameric PRMT5–CDK4–WDR77 complex (c.f. Fig. 2E, bottom left).

**Figure S3:**
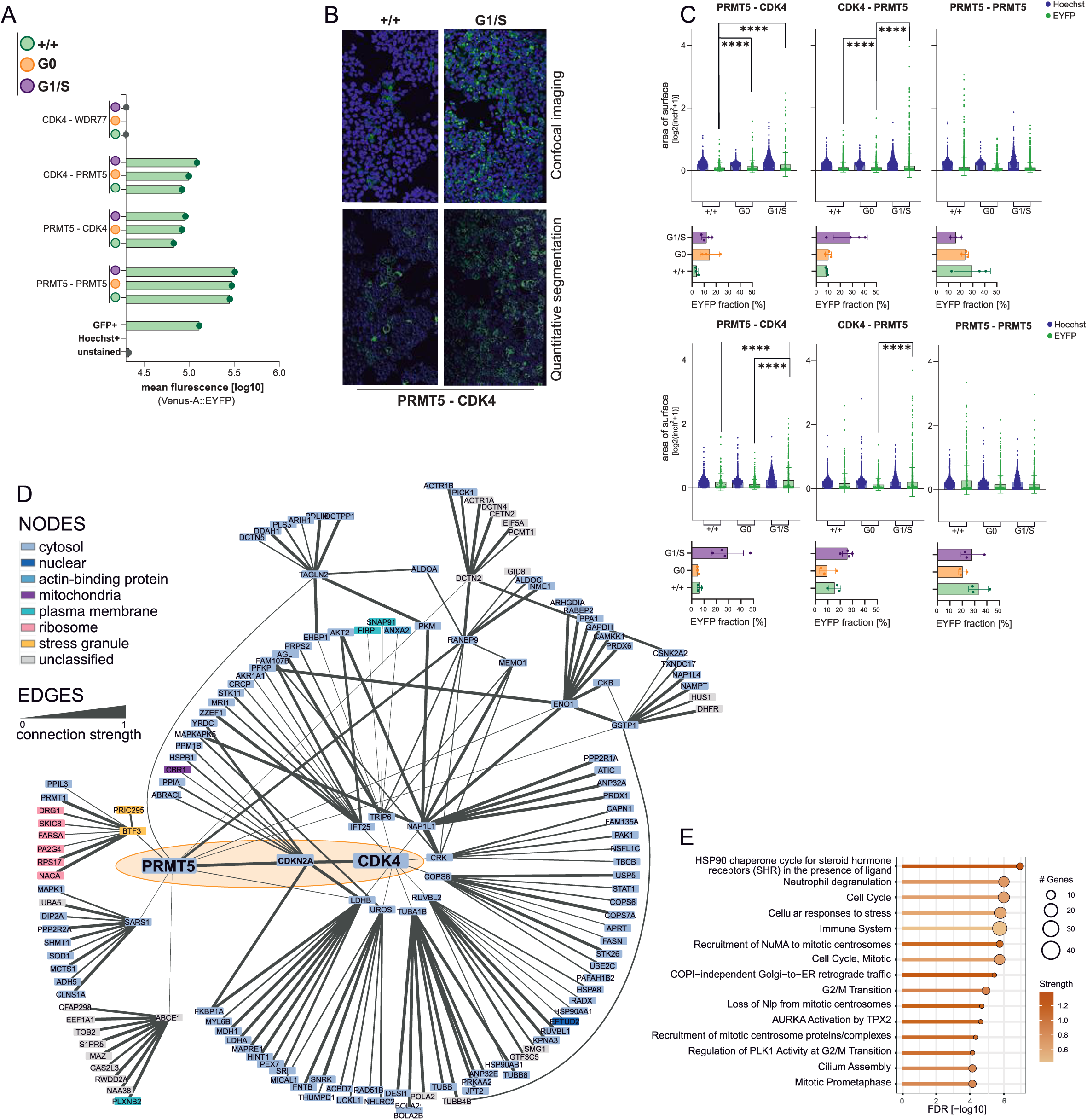
PRMT5 interacts with CDK4 in the cytosol in a cell cycle phase-dependent manner, independent of its methyltransferase activity. **(A)** Flow cytometry analysis of EYFP-positive cells across tested protein pairs and treatment conditions. Mean fluorescence intensities (log_10_) in the Venus-A::EYFP channel are shown. **(B)** Representative quantitative image analysis of PRMT5–CDK4 EYFP complementation in 293T cells under distinct cell cycle arrest conditions. Surface areas were measured for EYFP (interaction signal) and Hoechst33342 (nuclear DNA content), with merged images reflecting EYFP signal normalized to cell content. **(C)** Quantification of a second (top) and third (bottom) independently performed experiments highlighting the reproducible accumulation of PRMT5-CDK4 interaction increases during a G1/S-dependent cell cycle block. Area of surfaces [log2(inch^2^+1)] within the indicated channels and the resulting EYFP fraction [%] are summarized in a bar graph. (Significance of upregulated EYFP-signal obtained via two-tailed Wilcoxon test with p-values adjusted for multiple testing using FDR correction * < 0.05, ** < 0.01, *** < 0.001, and **** < 0.0001). (**D**) Spatial protein networks of PRMT5 and CDK4 derived from Organelle-IP data, with node color denoting subcellular compartment and edge thickness reflecting localization similarity (as described in Fig. 3E). (**E**) Functional enrichment analysis of unique PRMT5 and CDK4 network neighbors from (**D**), showing the top 15 Reactome pathways ranked by FDR using String-DB analysis.

**Figure S4:**
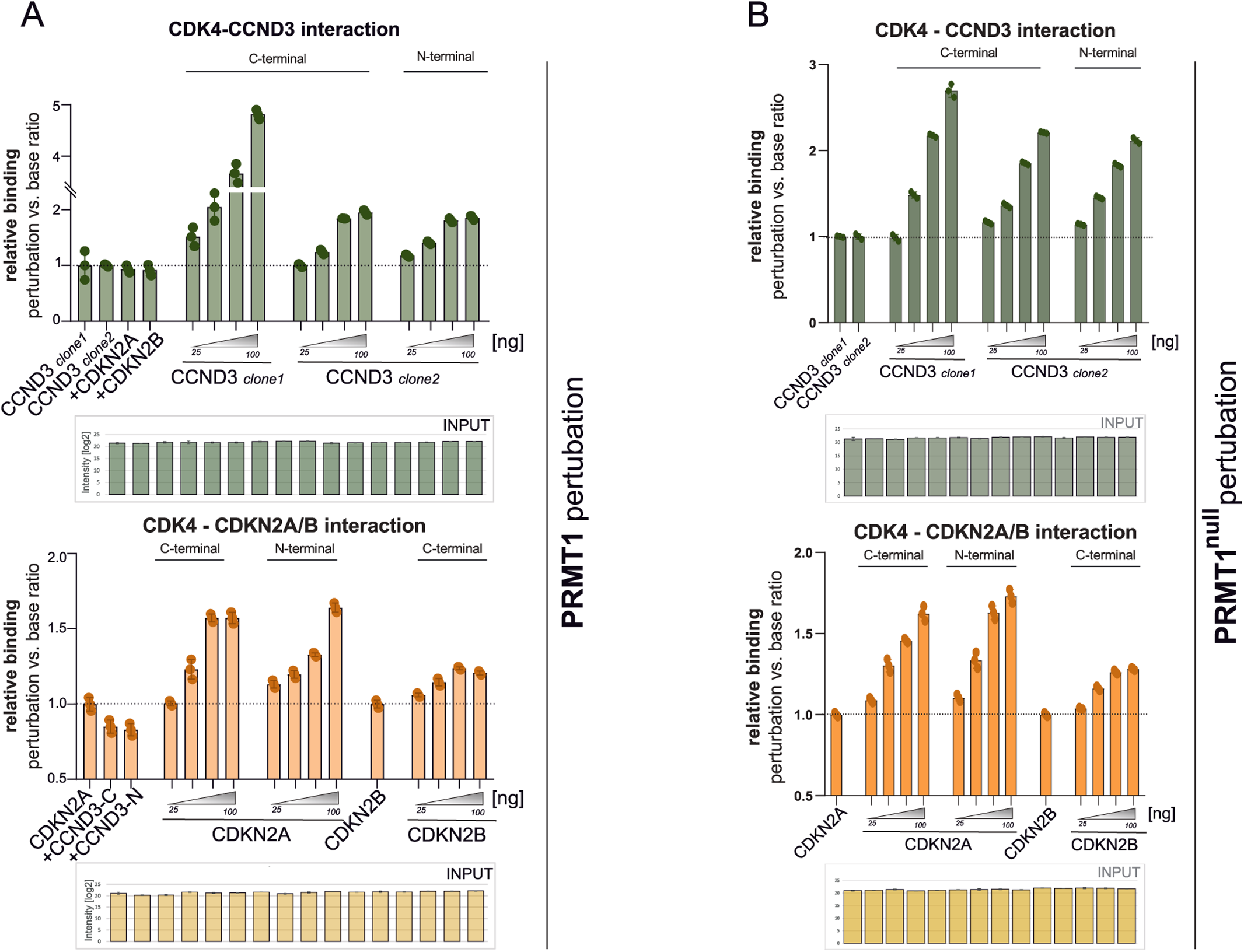
PRMT5 increases formation of the CDK4-CCND and CDK4-CDKN2A complexes independently of its methyltransferase activity. (**A**) Relative binding ratios from NanoLuc-based co-immunoprecipitation assays assessing the effect of PRMT1^WT^ perturbation on CDK4 interactions with CCND3 (green) and CDKN2A/B (orange). Data represent triplicates from three independent experiments, normalized to unperturbed controls. (**B**) Same analysis as (G), using the catalytically deficient PRMT1^null^ mutant (W215L).

**Figure S5:**
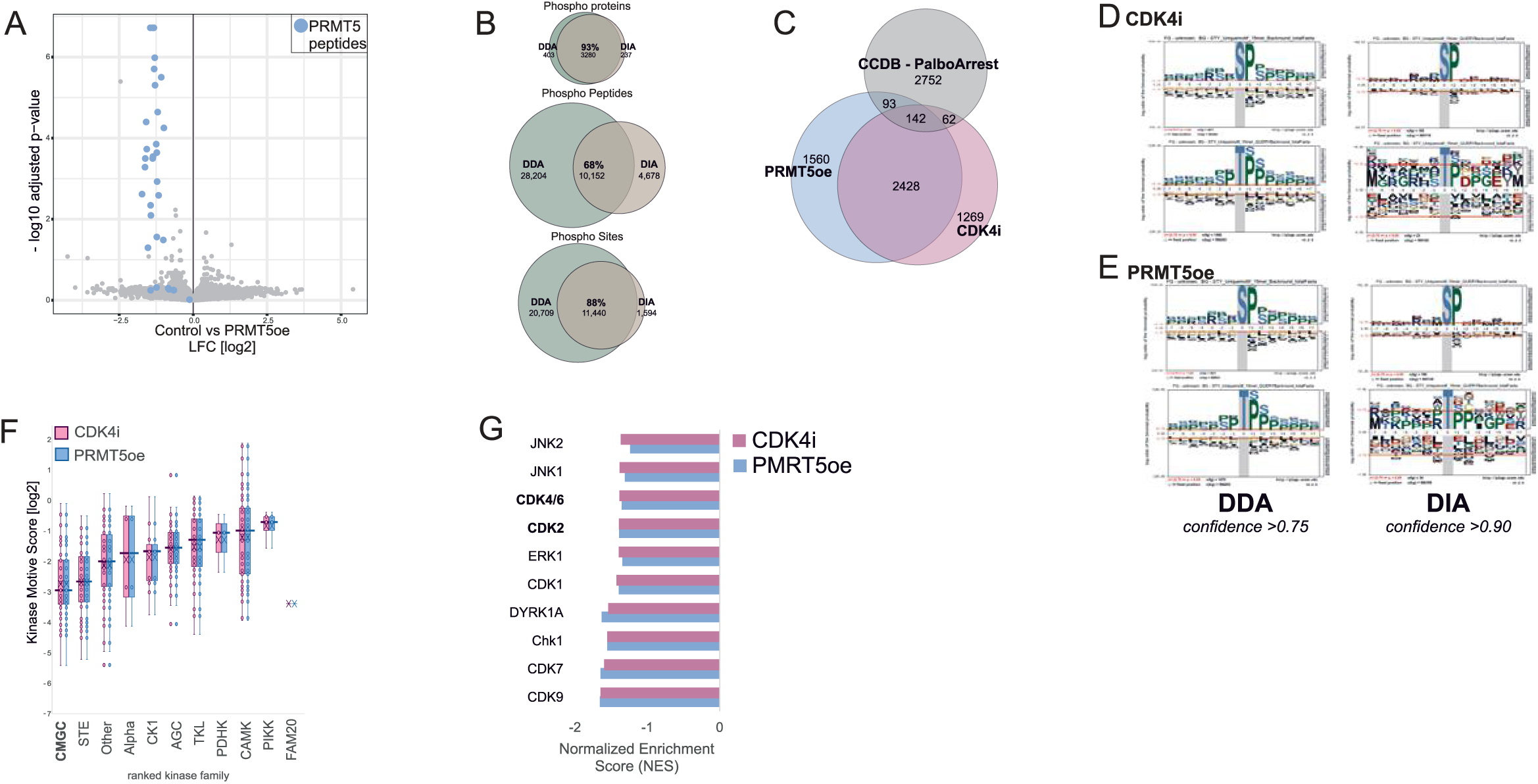
PRMT5 overexpression phenocopies/mimics CDK4 inhibition, with strongest phospho-suppression in RNA processing and cell cycle pathway. (**A**) Volcano plot of input samples comparing PRMT5oe vs. control, highlighting significantly altered peptides. PRMT5 peptides shown in light blue. (**B**) Venn Diagram displaying the phospho-protein (top), phospho-peptides, and phospho-site (bottom) overlaps between the different MS acquisition methods. (**C**) Venn diagram showing overlap of downregulated phospho-sites (log2FC ≤ –1.5) between CDK4i and PRMT5oe conditions, including reference overlap with palbociclib-responsive sites from the Cell Cycle Database (CCDB), related to Fig. 5F. (**D-E**) Probabilistic motif of the most downregulated phospho-sites (FG: log2FC < –1.5, BG: fasta-derived motives) for serine and threonine phospho-sites quantified in DDA (left) and DIA (right) upon CDK4i (D) (n(DDA-S/T) = 4211/1408; n(DIA-S/T) = 185/ 23) and PRMT5oe (E) (n(DDA-S/T) = 4371/ 1479; n(DIA-S/T) = 166/ 34). related to Fig.5I-J. (**F**) Boxplots from kinase motif analysis demonstrate a shared phospho-signature between PRMT5oe and CDK4i, with CMGC family kinases most strongly affected. (**G**) KSEA enrichment analysis of aggregated absolute log2FC values of downregulated (< –1.5) substrate phospho-sites of the indicated kinases within the CMGC family with their corresponding normalized enrichment score (NES) in the PRMT5oe and CDK4i condition, highlighting the top 10 hits with almost identical kinase substrate enrichments.

## Acknowledgements

We thank Christina Moesslacher, Johanna Kohlmayr, and Sandra Fasching for providing the yeast kinase-Y2H matrix. We thank Natalia Kunowska and Martina Derler for help in setting up the FACS experiments. We thank Natalia Kunowska for helping with phospho-proteomics protocol. We thank Sebastian Riedelbauch (Aarhus University) for providing access to the infrastructure for the AlphaFold multimer predictions. We thank Verina Manojlović and Hanna Engelke for help with the confocal microscopy setup and data quantification. The work was supported by BioTechMed-Graz through the Flagship project DYNIMO. This research was funded in whole, or in part, by the Austrian Science Fund (FWF) [10.55776/PAT8834124, 10.55776/P34316, 10.55776/COE14]. Mass spectrometry-based proteomics was supported by the Field of Excellence BioHealth - University of Graz.

## Author contributions

Conceptualization: SM, US

Data curation: SM, US

Formal Analysis: SM, BHW, US

Funding acquisition: US

Investigation: SM, TL, AEL, EA, MM

Methodology: SM, MM, US

Project administration: SM, US

Resources & Supervision: US

Validation: SM, TL, AEL

Visualization: SM, BHW, US

Writing – original draft: SM, US

Writing – review & editing: SM, US, and all authors

## Additional information

Supplementary information:

- Supplementary figures S1-S5
- Supplementary tables 1-8
- PDB files

